# Quantitative Evaluation of Normal Cerebrospinal fluid flow in Sylvian aqueduct and perivascular spaces of middle cerebral artery and circle of Willis using 2D phase-contrast MRI imaging

**DOI:** 10.1101/2025.02.11.637564

**Authors:** Rosemarie Faustina D. Le, Christof Karmonik, Angelique S. Regnier-Golanov, Eugene V. Golanov, Gavin W. Britz

**Affiliations:** Houston Methodist Academic Institute, 6560 Fannin str., Suite 944, Houston, TX, 77030

## Abstract

Recently, it was proposed that CSF flow constitutes a critical part of the glymphatic system, playing a critical role in various brain abnormalities from Alzheimer’s disease to hydrocephalus. Thus, measurement of CSF flow has been increasingly used for diagnostic and clinical monitoring purposes. Phase-contrast MRI has been used to determine CSF flow. However, CSF flow in the periarterial spaces of the circle of Willis and the middle cerebral artery which are important conduits remain unexplored. We employed phase-contrast MRI to explore CSF flow along the perivascular spaces of the circle of Willis and the middle cerebral artery to establish baseline parameters of CSF and compare them with the Sylvian aqueduct. To analyze CSF flow in the perivascular space, we developed a new, semi-automated method for outlining the perivascular space and extracting CSF flow parameters. The twenty-four healthy participants were recruited to achieve an even distribution by age (mean: 40 ± 11) and gender (13 males). We validated our routine for CSF flow measurements by comparing CSF flow in the Sylvian aqueduct (0.00700 mL/s) with the range of literature values, 0.0049-0.0432 mL/s. For all CSF parameters, the circle of Willis and middle cerebral artery were differed from the Sylvian aqueduct. For most CSF flow parameters, the 95% confidence intervals of the circle of Willis and middle cerebral artery overlap. The linear mixed models and general linear mixed models for flow indicate strong effects of the conduits. CSF velocity in these conduits were lower 0.159 cm/s and 0.198 cm/s respectively than in the Sylvian aqueduct. Overall, differences in CSF flow parameters between sex and age groups were negligible. In this study, we have validated our routine and established baseline values of CSF flow along the circle of Willis and the middle cerebral artery as well as highlighted the limited influence of sex and/or age.

## Background

Cerebrospinal fluid (CSF) is a clear fluid surrounding the central nervous system (CNS): brain, filling cerebral ventricles, and central canal of the spinal cord. At any given time, there are 90-150 mL of CSF in the cranium and spinal cord because production and absorption occur at the same rates to maintain intracranial pressure (1, 2). CSF is produced in the choroid plexus of the lateral ventricles and flows through the interventricular foramina to the third ventricle, along the Sylvian aqueduct (SA) to the fourth ventricle, proceeding to the subarachnoid space, and to the spinal cord (1–3). Recently, the concept of the glymphatic pathway has been introduced (4, 5). According to it CSF flows from the subarachnoid space to periarterial space and enters the brain parenchyma where it mixes with the interstitial fluid and further exits parenchyma into the paravenous space. CSF flow is pulsatile due to cardiac pulsations and breathing (3, 6). CSF moves in opposite directions depending on the cardiac cycle: during systole, intracranial arteries increase in volume which pushes CSF in the craniocaudal direction. The reverse happens in diastole as blood vessel volume decreases and CSF moves in the caudocranial direction (2, 7, 8).

CSF serves several physiologic functions. CSF provides support and cushioning for the brain. Movement of CSF through the brain parenchyma clears out waste and transport essential molecules (1–3, 9, 10). These functions and dynamics, especially CSF flow and pressure, can be altered in pathologies. As a part of the glymphatic system, CSF flow has been proposed to play a role in Alzheimer’s disease by failing to clear out amyloid beta and tau tangles, the characteristic protein markers (3, 11, 12). Abnormal increase in intracranial pressure due to abnormal CSF drainage is associated with hydrocephalus (7, 8, 11, 12). Other studies have observed the roles of CSF flow in meningitis, cerebral edema, and other cerebrovascular diseases (8, 12). Thus, measurement of CSF flow has been increasingly used for diagnostic and clinical monitoring purposes (13).

To explore the clinical implications of CSF flow, many studies focused on the CSF flow through the SA which connects the third and fourth ventricles (1, 10). SA containing only CSF flow is easier to identify and image, so it frequently has been studied. On the other hand, the periarterial spaces of circle of Willis (COW) and the middle cerebral artery (MCA), important conduits of CSF flow, remain unexplored. The COW localized to the base of the brain formed by anterior cerebral arteries, anterior communicating artery, internal carotid arteries (at their distal tips), posterior cerebral arteries, and posterior communicating artery. MCA originates from COW at the internal carotid arteries joining and ascending in the lateral sulcus of the cerebrum providing blood supply to many parts of the lateral cortex.

Therefore, this study has several objectives. First, because the CSF flow along the perivascular spaces of COW and MCA have not been studied yet, we aim to establish baseline measurements of CSF flow parameters along these conduits, including potential differences due to sex and age. Second, we aim to compare CSF flow parameters between the SA, COW, and MCA.

For this study, we used phase-contrast magnetic resonance imaging (PC-MRI). PC-MRI was first used in 1994 by Barkhof et al. (14) to evaluate the effect of age on SA flow (12). Since then, PC-MRI has emerged as the gold standard for measuring CSF flow (15). The advantages of PC-MRI are that it is noninvasive and relatively quick at taking measurements (12).

To analyze CSF flow in the perivascular space, we developed a new, semi-automated method for outlining perivascular space and extracting CSF flow and velocity parameters (Additional file 1). Many semi-automated or automated methods have been developed in order to increase accuracy, save time on analysis, and/or contribute to study reproducibility (16, 17). We conducted preliminary evaluation of our semi-automated program, through validation with the CSF flow through SA parameters which are extensively covered in the literature.

We focused on the following CSF flow parameters: stroke volume (StroVol), volumetric flow rate (VFR), systolic flow rate (SFR), diastolic flow rate (DFR), velocity, peak systolic velocity (PSV), and peak diastolic velocity (PDV). While the most useful parameters have yet to be determined, we chose these parameters to capture the overall picture of CSF flow dynamics in the hopes that they may be used for future clinical diagnostics and therapeutics (18).

## Methods

### Participants

The twenty-four healthy participants were recruited to achieve an even distribution by age and gender (13 males). Ages ranged from 23 to 59 with a mean of 40 ± 11 (Table 1). To obtain baseline parameters by demographics, participants were separated into different age groups: 20-29, 30-39, 40-49, and 50-59. The study was approved by the Houston Methodist Hospital Institutional Review Board, protocol Pro00022145.

**Table 1.** Participant demographics.

| *Demographics* | *Description* | *Number* | *Percentage* |
| --- | --- | --- | --- |
| Age | 20-29 | 6 | 25% |
|  | 30-39 | 4 | 17% |
|  | 40-49 | 7 | 29% |
|  | 50-59 | 7 | 29% |
|  | Total | 24 | 100% |
| Sex | Male | 13 | 54% |
|  | Female | 11 | 46% |
|  | Total | 24 | 100% |

### MRI Acquisition

A 2D PC-MRI image was acquired at each region of interest (COW, MCA, SA) chosen by an experienced neuroscientist (EG, ARG). Details of the acquisition parameters are as follows: slice thickness 3 mm, acquisition matrix 272 x 272, FOV 160 cm x 160 cm, in-plane resolution 0.6 mm x 0.6 mm. TE=3 msec, TR=105 msec. Images were acquired with a single transmit, 32 channel receive head coil in the FDA-approved clinical mode of the MAGNETOM 7 Tesla human MRI scanner (Siemens Healthineers, Erlangen, Germany). A 3D velocity encoding scheme was used to account for CSF flow direction not exactly parallel to the slice normal. The encoding velocity (VENC) was set at 5 cm/s to remain sensitive to the corresponding low velocity of CSF flow and to eliminate blood flow contamination which can be recognized as global maximum/minimum gray scale values due to the phase wrapping artifact of blood velocities exceed this VENC value. A total of 20 images evenly spaced across the cardiac cycle were reconstructed with a total acquisition time of approximately 5 min for each location.

### CSF Flow Analysis

As the VENC value was low compared to blood flow velocities, global maximum black/white gray scale intensity values –-caused by the phase wrapping artifact-- were used as a mask for distinguishing and eliminating blood flow from perivascular CSF flow in 2D phase contrast images. Regions of interest (ROI) in an annular shape centered on the arterial cross section were chosen for quantifying CSF flow using a semi-automated algorithm described in Additional file 1.

Magnitude images were used as anatomical references. Two ROIs --one left and one right-- were identified for each COW and MCA and one ROI was identified for each SA (Fig. 1). The established ROI settings were entered into our developed in-house semi-automated program to produce flow rate (FR) and Velocity values. FR (mL/s) was calculated as the area of the ROI (converted from mm^2^ to cm^2^) x VENC x (mean intensity / maximum gray scale intensity). Velocity (cm/s) was calculated as FR / the area of the ROI. ROIs for 20 cardiac gated images over one cardiac cycle were obtained from each series to obtain the FR and Velocity curves.

**Figure 1.**
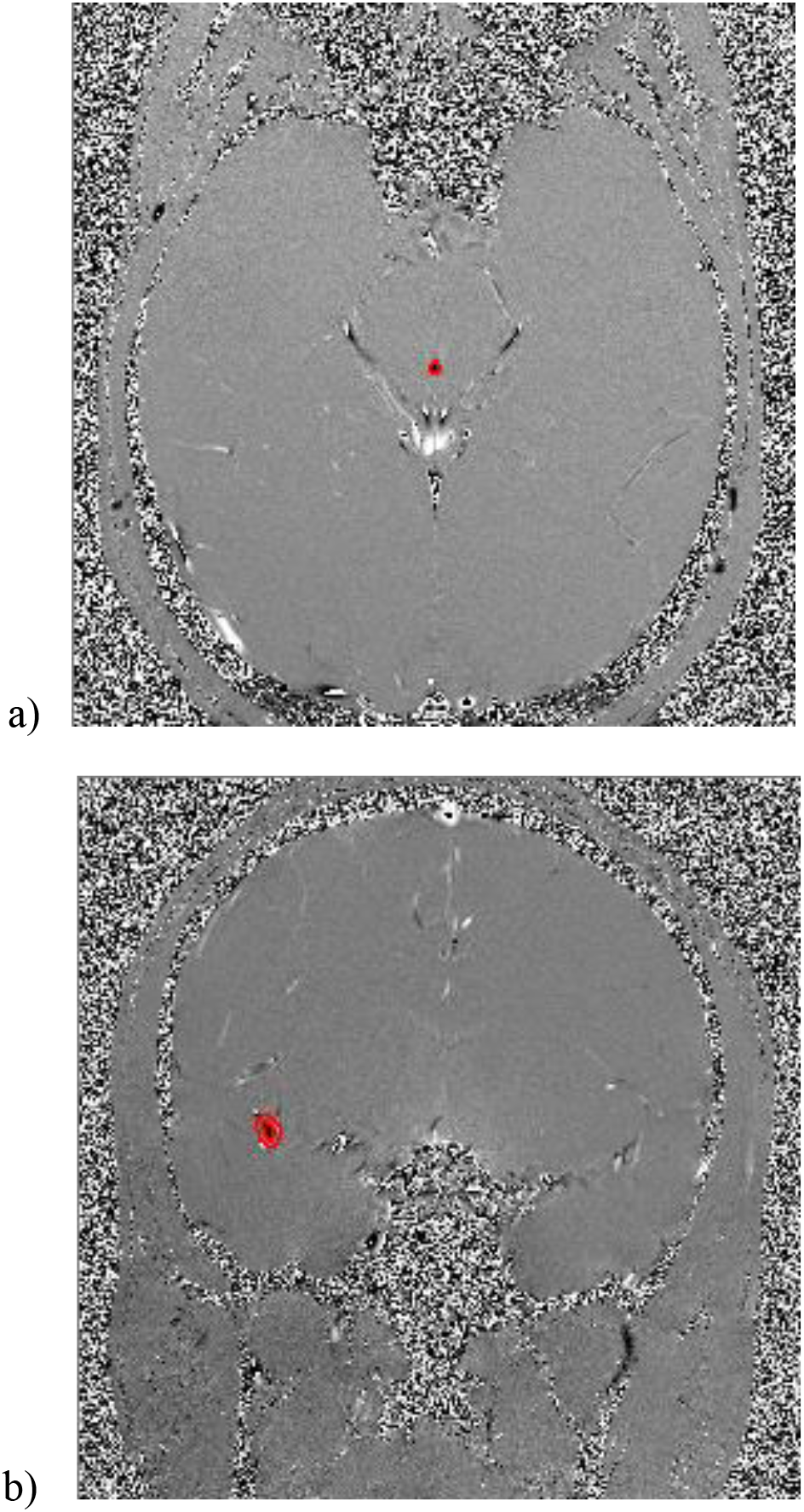
Examples of ROIs. (a) ROI of SA and (b) one of two ROIs of MCA on phase images. For the first phase image of the 20 series of images for the cardiac cycle, the X,Y-coordinates, tolerance, and band size of the ROIs were determined by the user (Additional file 1).

Peak timing was standardized by looking at which percentage of the cardiac cycle peak flow occurred. The percentage of the cardiac cycle was calculated as (the ordinal position of the image within the series / 20) x 100. The stroke volume was calculated as FR x time.

### Statistical Analysis

For all parameters in each location of SA, COW, and MCA, median values and bias-corrected and accelerated bootstrap (BCa) confidence intervals (CIs) from non-parametric bootstrapping were calculated.

To compare the conduits and evaluate the influence of demographics, various models were constructed. For StroVol, VFR, and Velocity, linear mixed models (LMM) were constructed with block bootstrapping. Generalized linear mixed models (GLMM) with the gamma family and log link function were constructed with block bootstrapping for SFR, DFR, and Area. Generalized linear models (GLM) were constructed for PSV and PDV. For these LMMs, GLMMs, and GLMs, BCa CIs were calculated. For time to peak as percentage of cardiac cycle duration (TP) of StroVol (TPStroVol), VFR (TPVFR), and Velocity (TPVelocity), beta regression with the logit link function was constructed, and Wald CIs were calculated. For comparing SA, COW, and MCA, fixed effects were the intercept (= SA), time (except for PSV, PDV, and TP parameters), COW, and MCA. SA was set as the intercept because CSF flow parameters for the SA are already established in the literature. For evaluating the influence of sex and age, fixed effects were the intercept, time (except for PSV, PDV, and TP parameters), sex, and age. An interaction term between sex and age was also added. When applicable, participants were considered as random effects to account for the non-independence of time series. Age was rescaled, zero random effects were dropped, and/or optimizers were changed to address model convergence issues as needed. Whenever the data contained zero-values, the following transformation was applied to the outcome variable before beta regression: (y x 23 + 0.5) / 23. For post-hoc testing, p-values were adjusted using the Benjamini-Hochberg procedure. Assumptions were checked with QQ plots and residual plots. All statistical analysis and visualization were performed in R/RStudio.

## Results

Baseline values of CSF flow parameters with the median and 95% CI by conduits and by sex and age are reported in Table 2 and Tables 3 and 4 respectively. Model estimates of the fixed and random effects are reported in Table 5 for conduits comparison and Table 6 for the effect of sex and age . StroVol is reported in mL. VFR, SFR, and DFR are reported in mL/s. Velocity, PSV, and PDV are reported in cm/s. TPStroVol, TPVFR, and TPVelocity are reported in percentage of the cardiac cycle. Area is reported in cm^2^.

**Table 2.** Median values of CSF flow parameters in SA, COW, and MCA.

|  | *Conduit* | *Median* | *Lower 95% CI* | *Upper 95% CI* |
| --- | --- | --- | --- | --- |
| StroVol (mL) | SA | 0.00254 | 0.00221 | 0.00277 |
|  | COW | 0.0146 | 0.0140 | 0.0156 |
|  | MCA | 0.0140 | 0.0131 | 0.0149 |
| VFR (mL/s) | SA | 0.00700 | 0.00600 | 0.00700 |
|  | COW | 0.0420 | 0.0400 | 0.0430 |
|  | MCA | 0.0450 | 0.0420 | 0.0460 |
| SFR (mL/s) | SA | 0.00900 | 0.00800 | 0.00900 |
|  | COW | 0.0530 | 0.0500 | 0.0570 |
|  | MCA | 0.0550 | 0.0530 | 0.0580 |
| DFR (mL/s) | SA | 0.00400 | 0.00400 | 0.00500 |
|  | COW | 0.0230 | 0.0220 | 0.0250 |
|  | MCA | 0.0270 | 0.0230 | 0.0280 |
| Velocity (cm/s) | SA | 0.578 | 0.578 | 0.723 |
|  | COW | 0.285 | 0.271 | 0.299 |
|  | MCA | 0.260 | 0.246 | 0.267 |
| PSV (cm/s) | SA | 1.45 | 1.16 | 1.45 |
|  | COW | 0.749 | 0.677 | 0.848 |
|  | MCA | 0.676 | 0.516 | 0.728 |
| PDV (cm/s) | SA | 0.867 | 0.578 | 0.867 |
|  | COW | 0.383 | 0.368 | 0.622 |
|  | MCA | 0.405 | 0.303 | 0.452 |
| TPStroVol (%) | SA | 50.0 | 40.0 | 62.5 |
|  | COW | 75.0 | 70.0 | 80.0 |
|  | MCA | 70.0 | 70.0 | 80.0 |
| TPVFR (%) | SA | 20.0 | 20.0 | 25.0 |
|  | COW | 40.0 | 30.0 | 55.0 |
|  | MCA | 40.0 | 15.0 | 42.5 |
| TPVelocity (%) | SA | 55.0 | 50.0 | 55.0 |
|  | COW | 65.0 | 50.0 | 65.0 |
|  | MCA | 65.0 | 55.0 | 75.0 |
| Area (cm<sup>2</sup>) | SA | 0.00692 | 0.00692 | 0.0104 |
|  | COW | 0.0277 | 0.0277 | 0.0311 |
|  | MCA | 0.0398 | 0.0381 | 0.0415 |

**Table 3.**
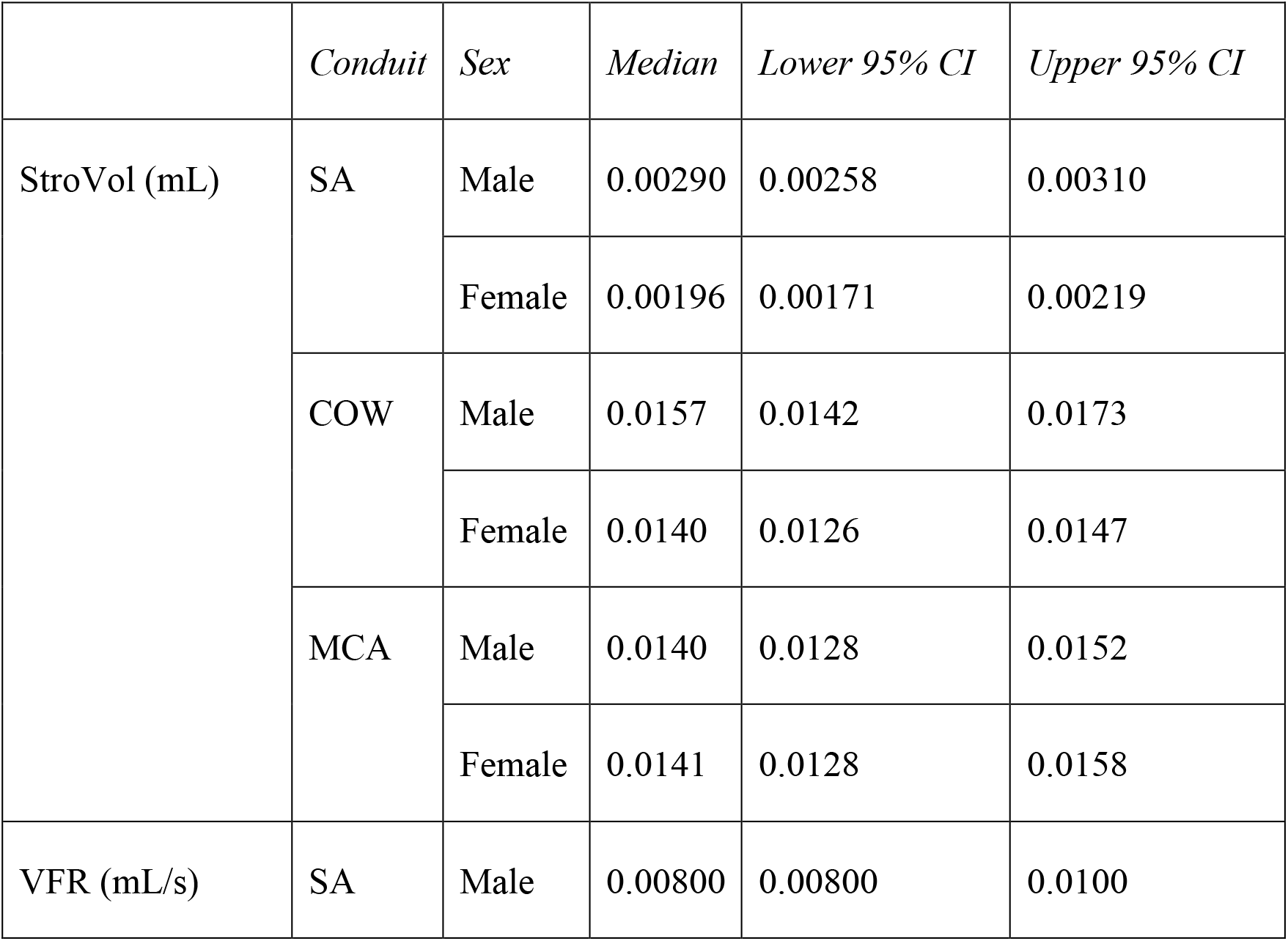
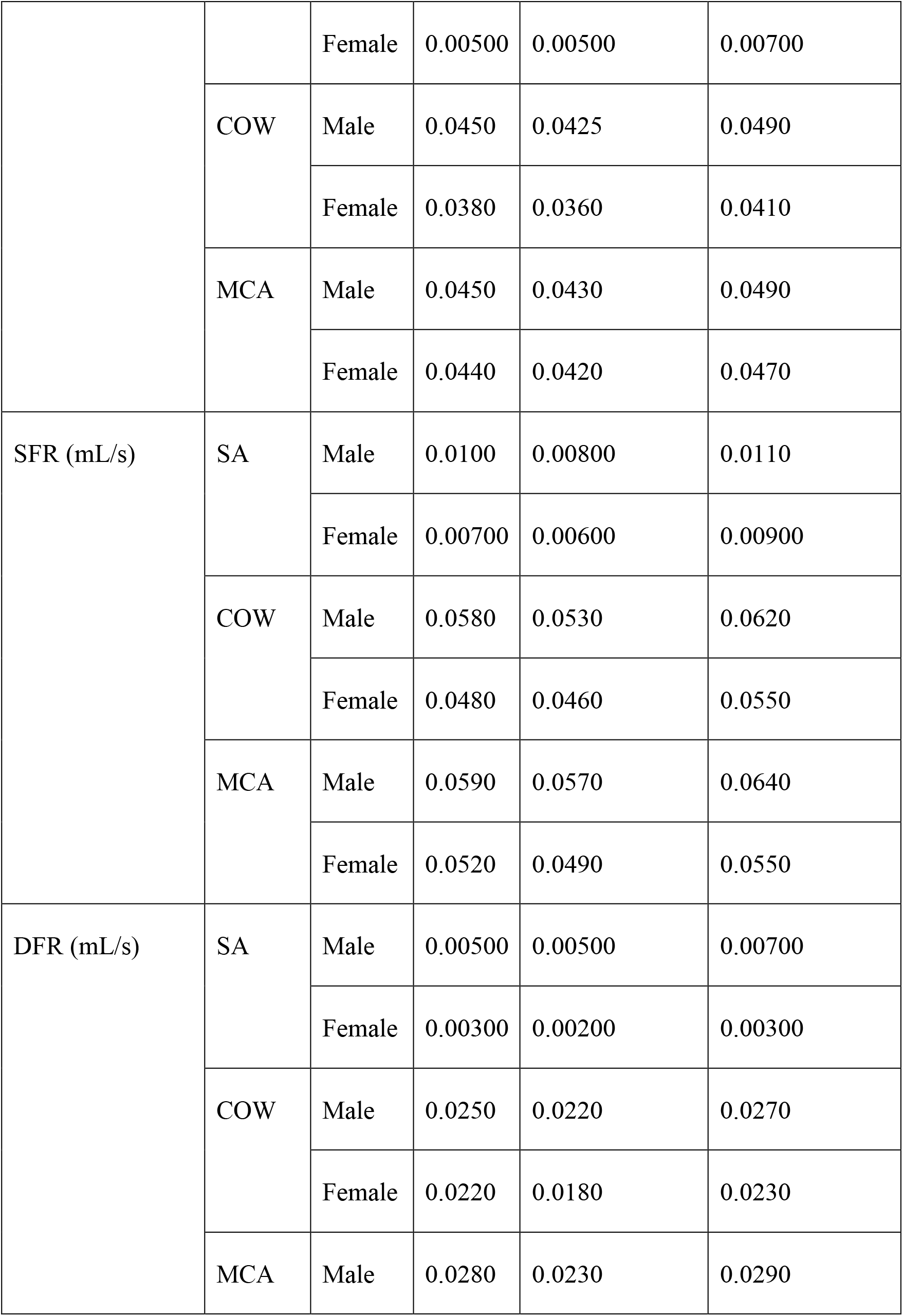
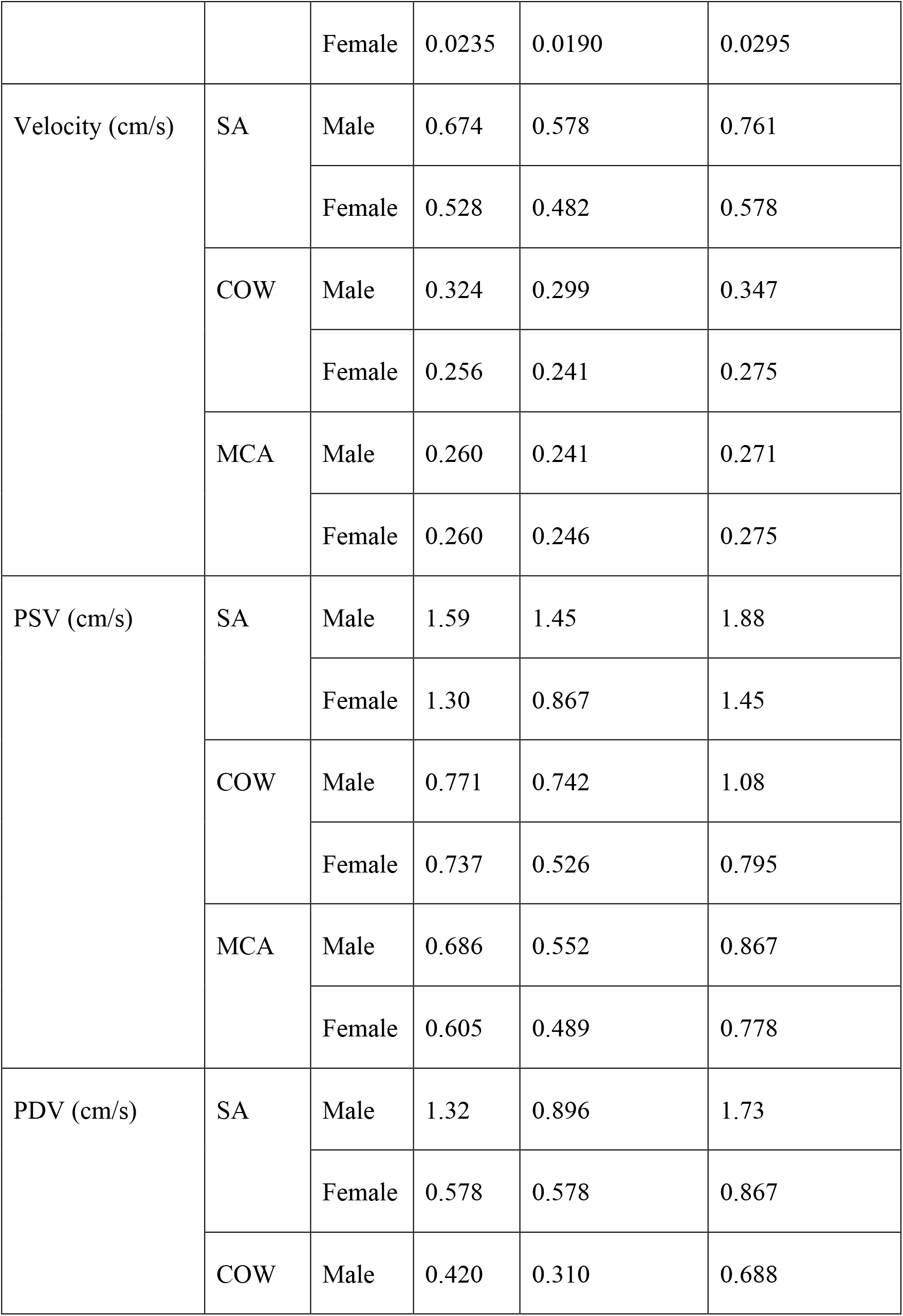
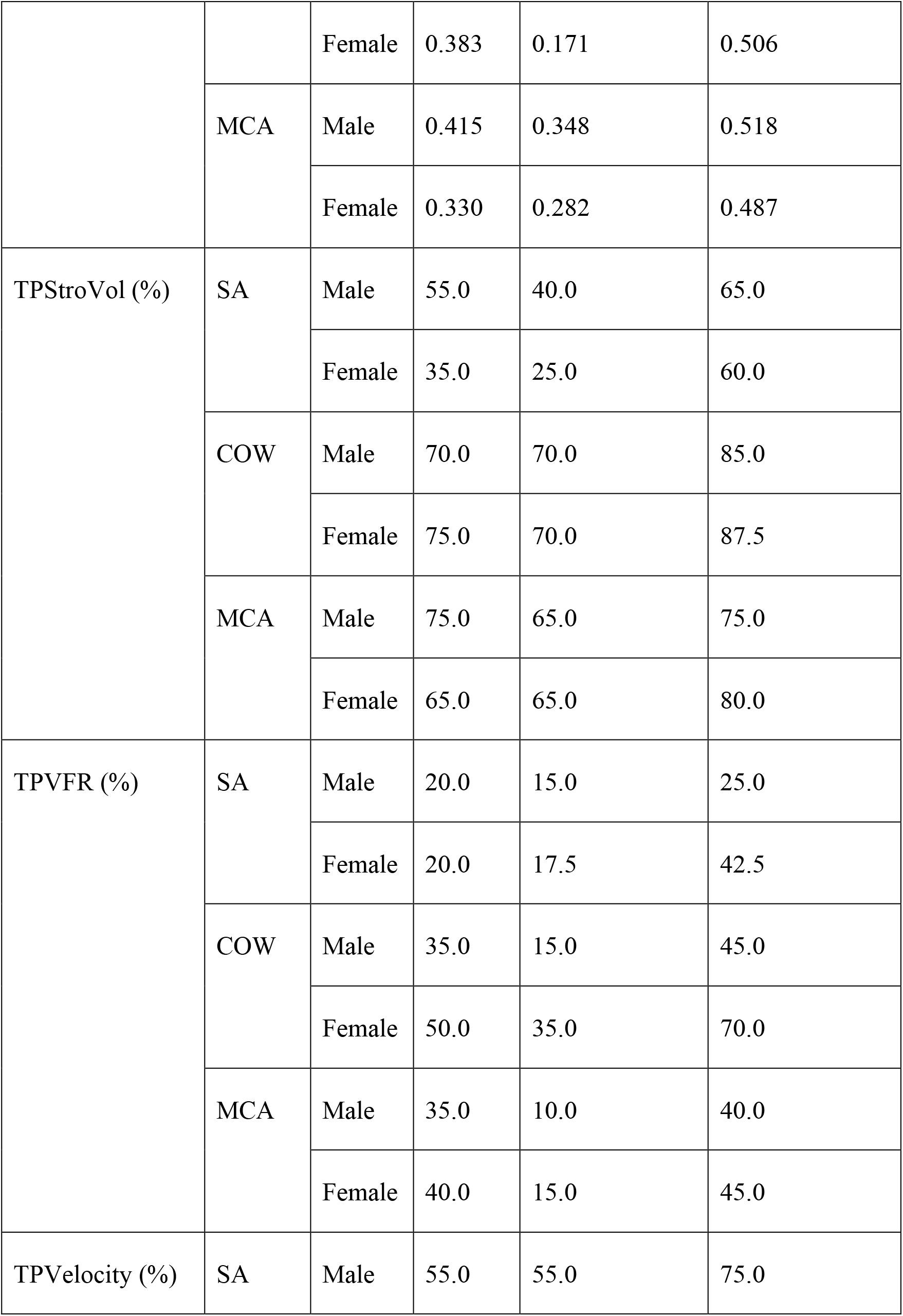
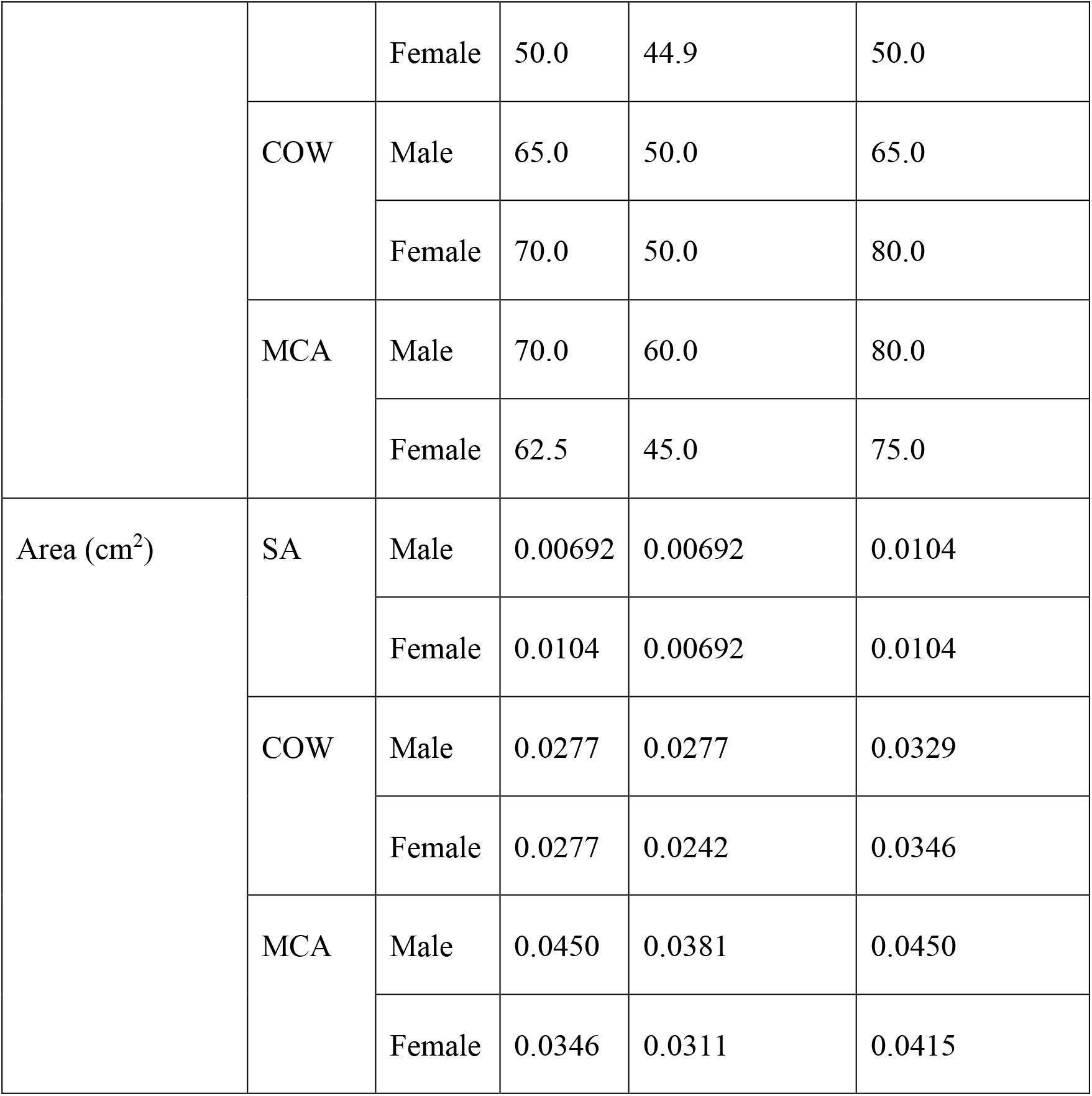
Median values of CSF flow parameters in different conduits by sex.

**Table 4.**
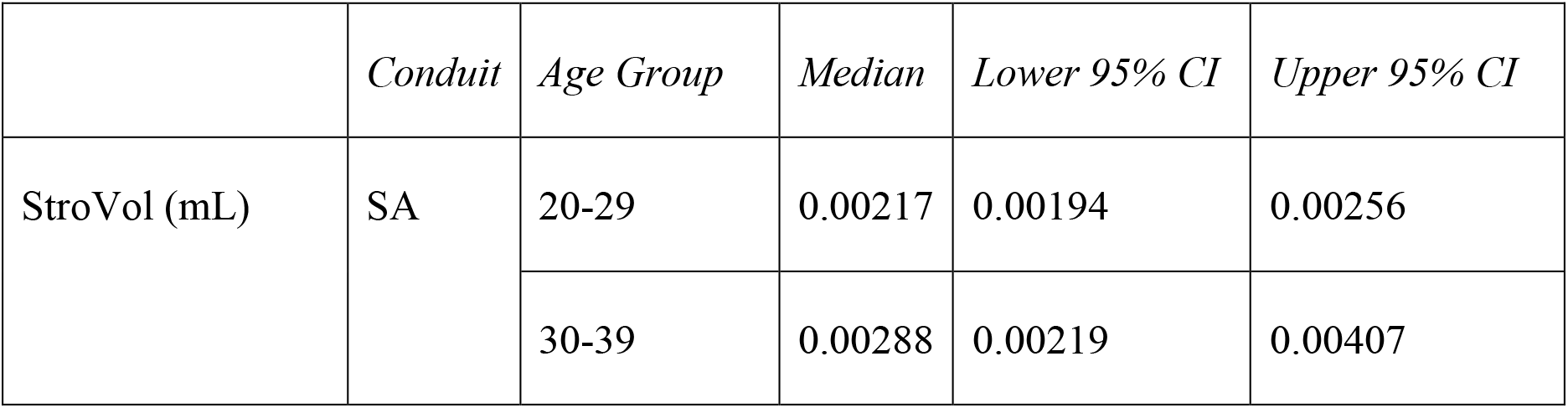
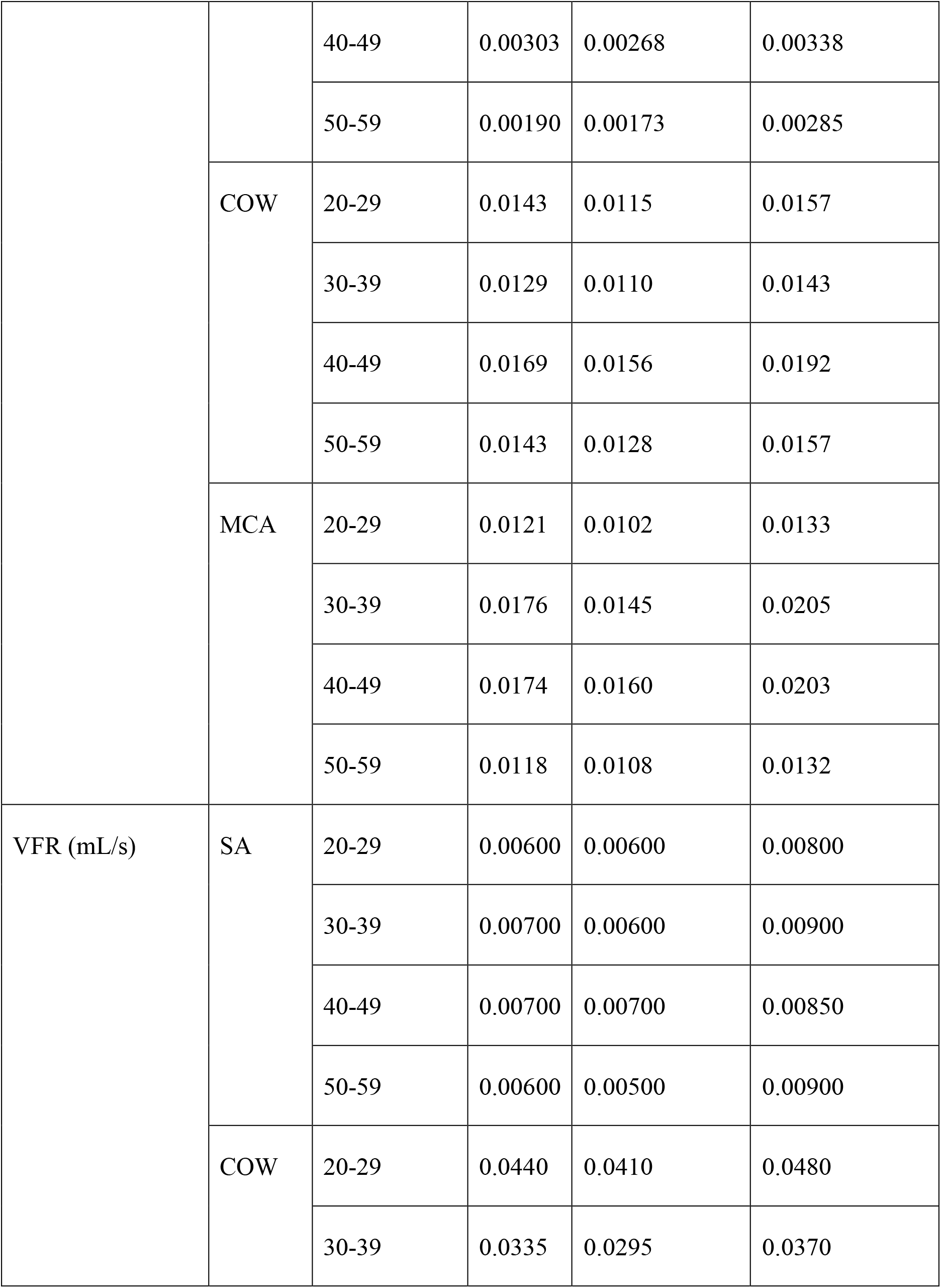
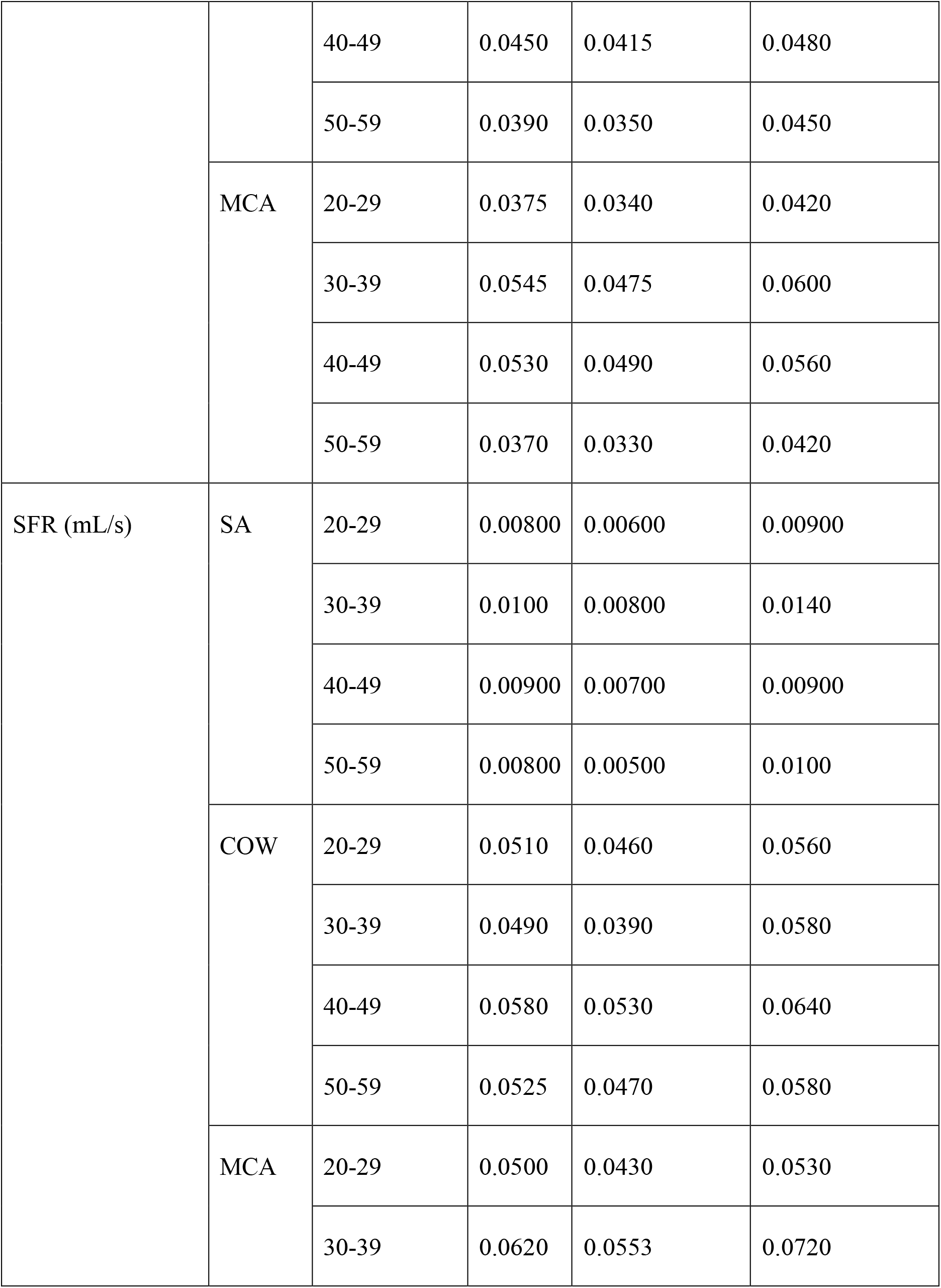
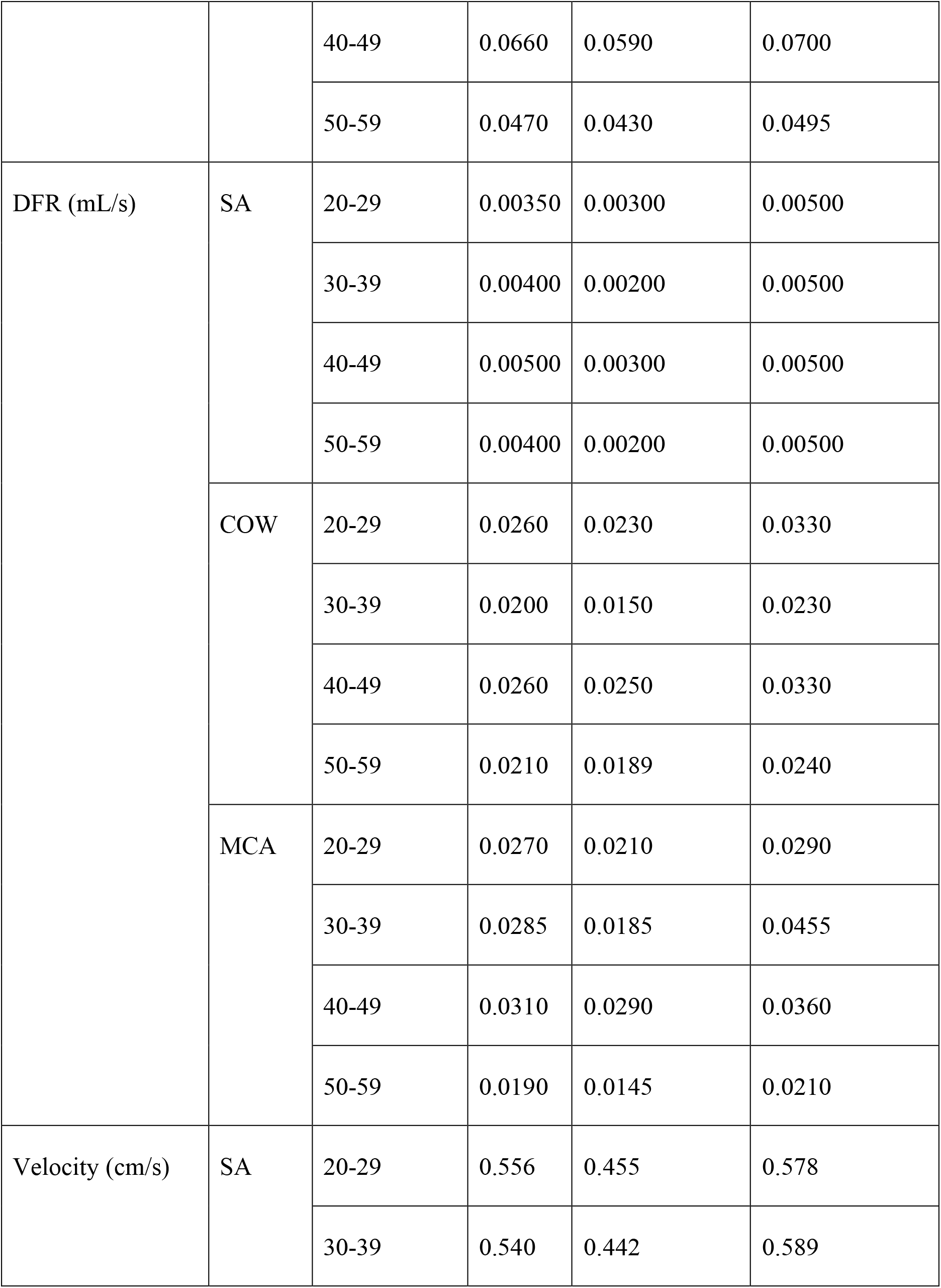
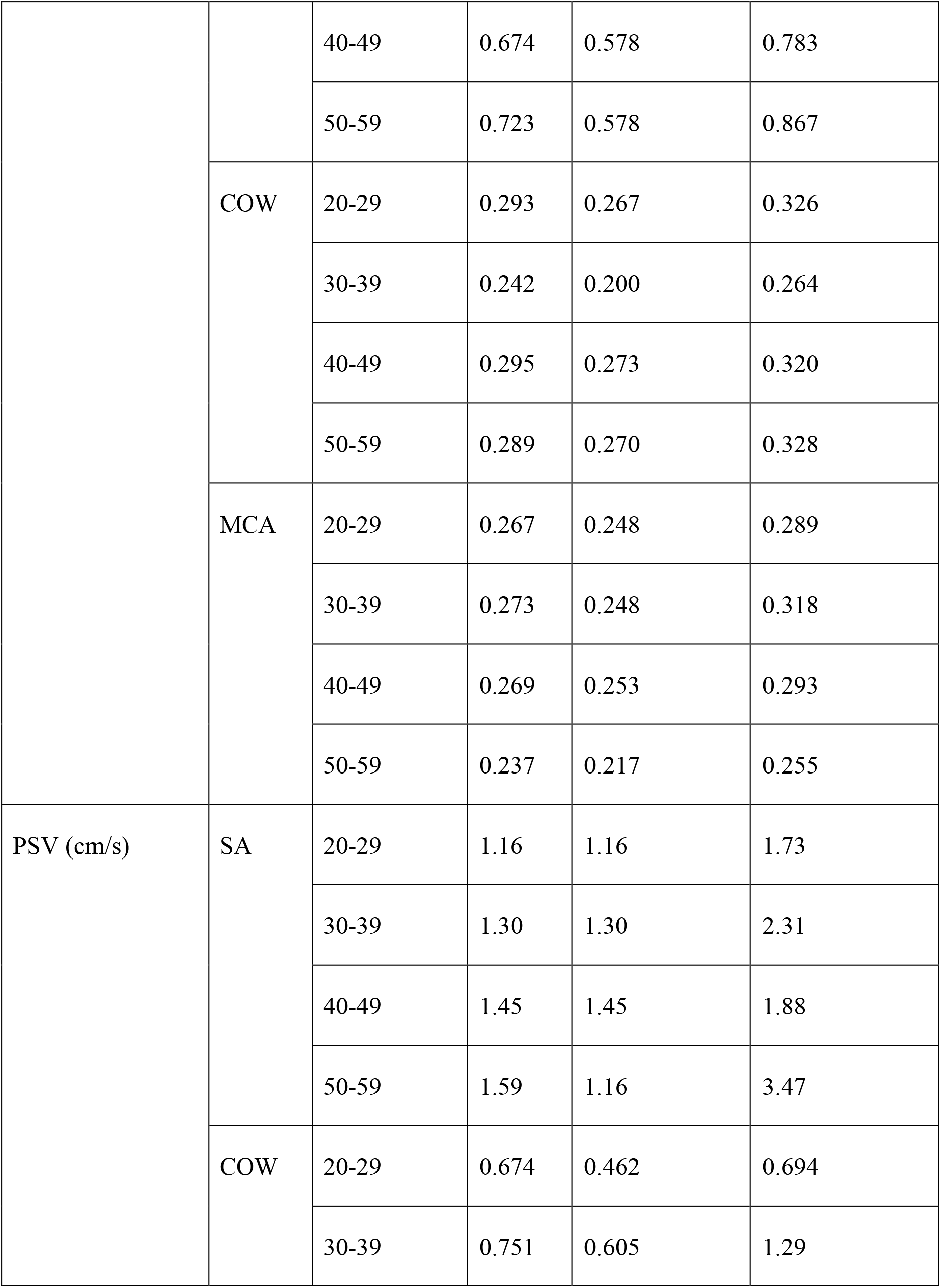
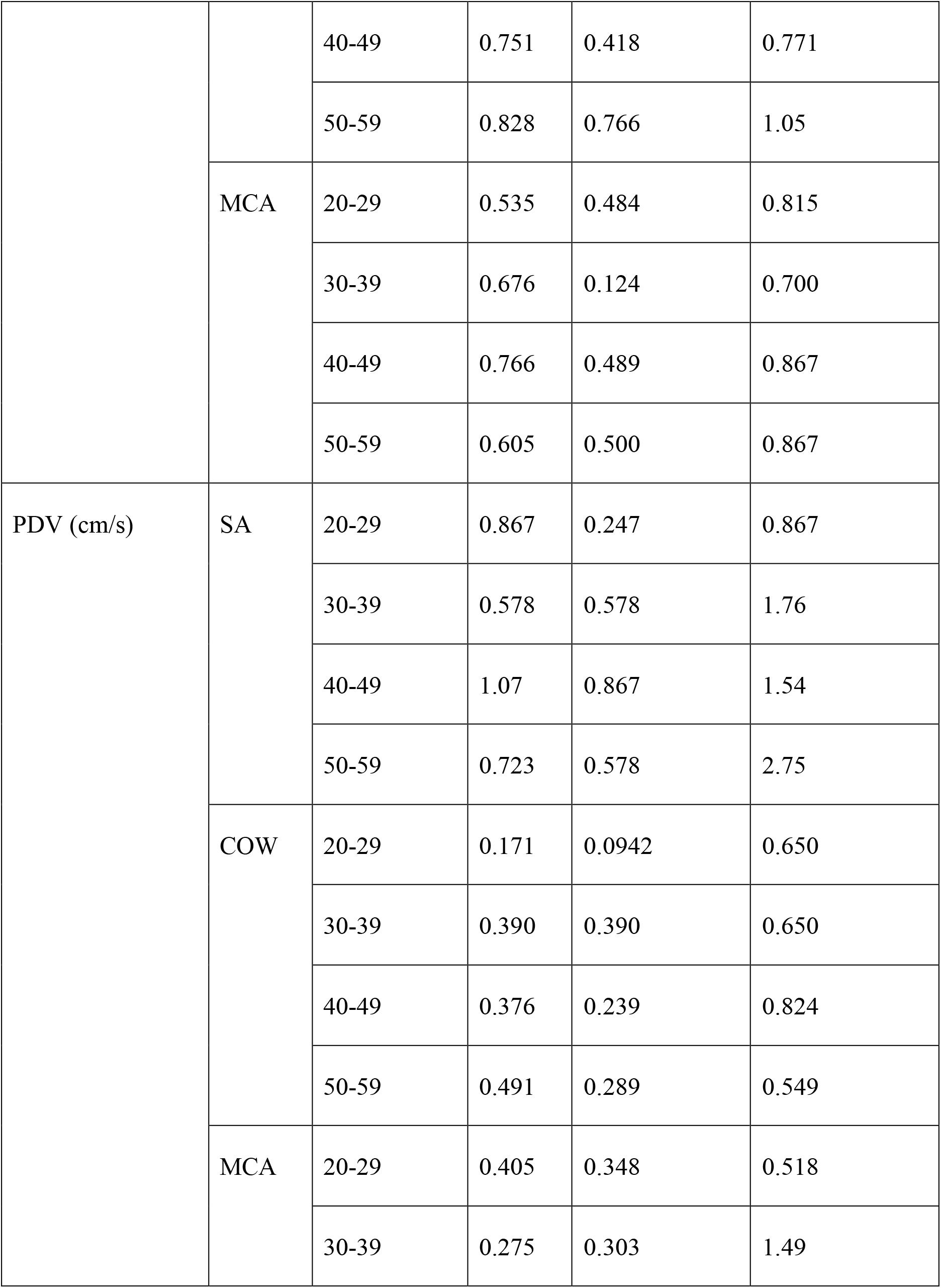
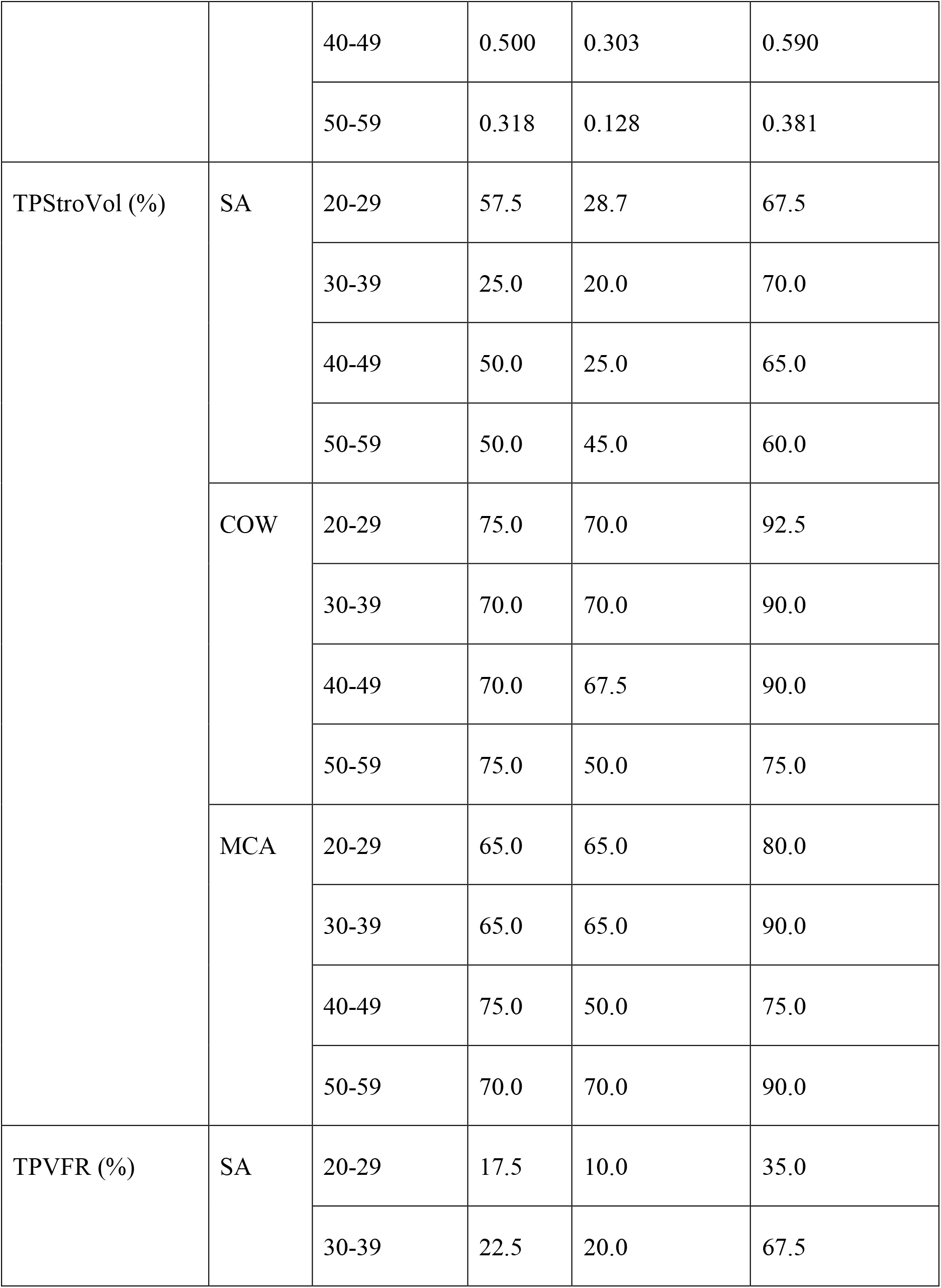
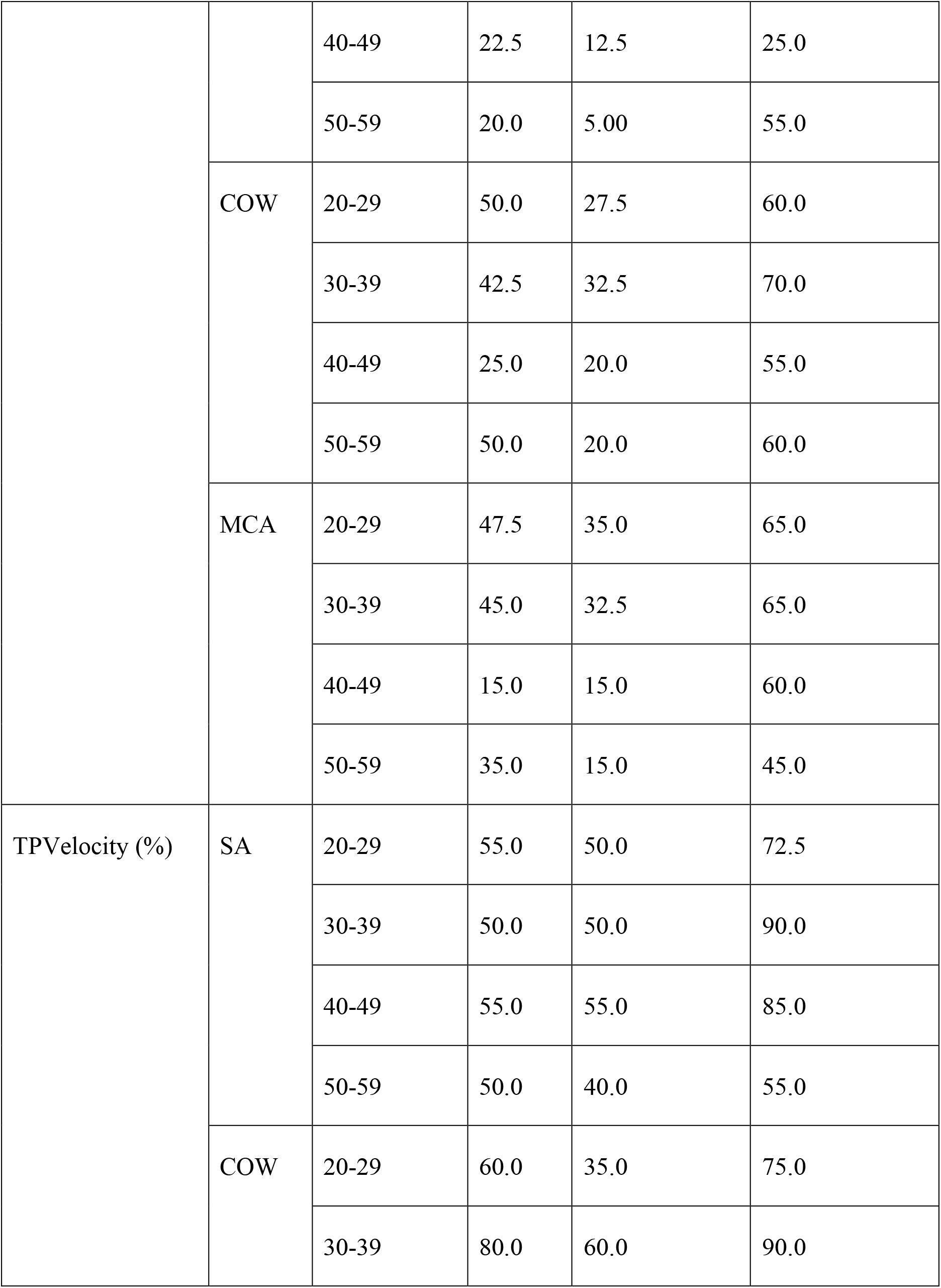
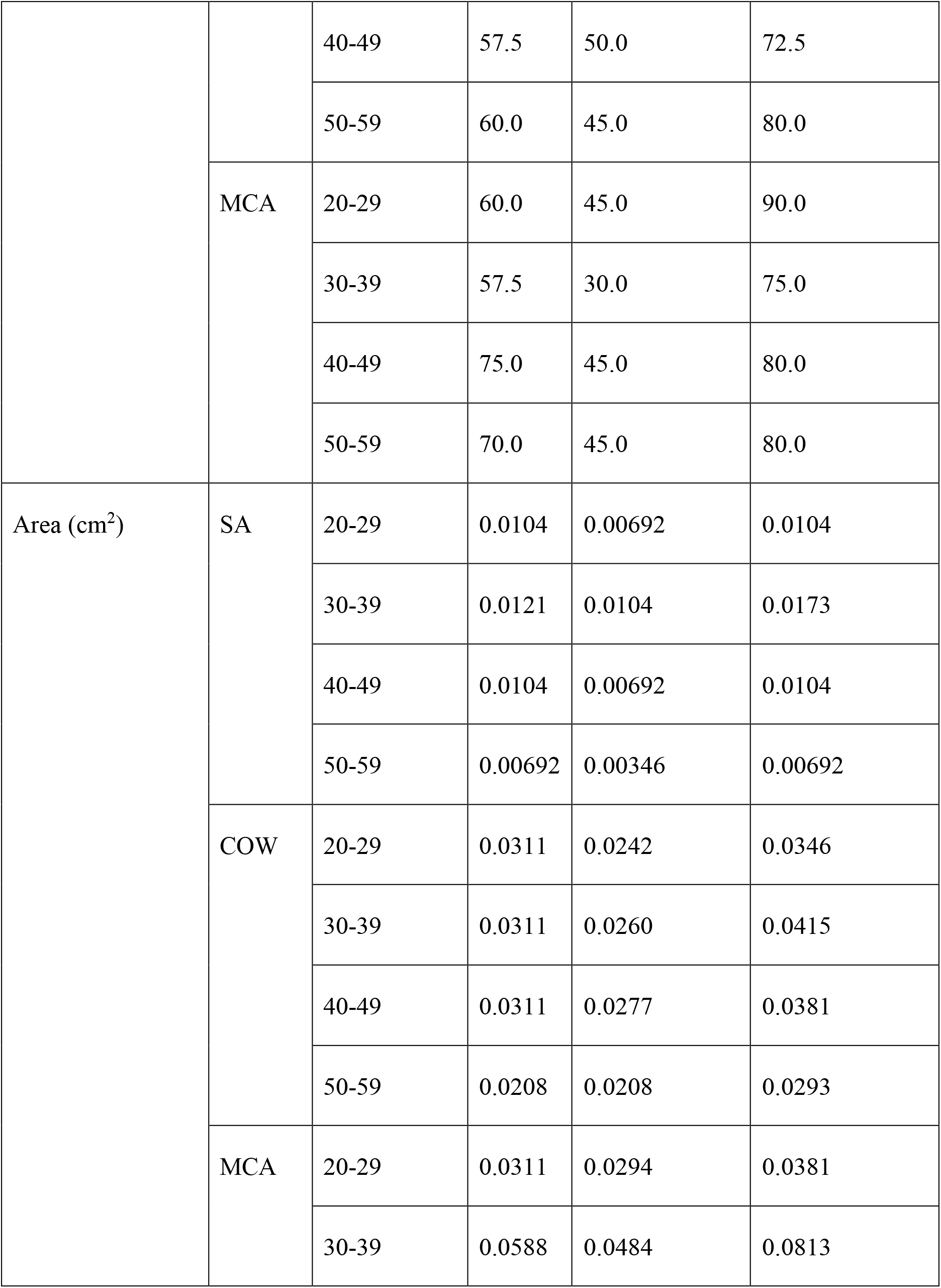
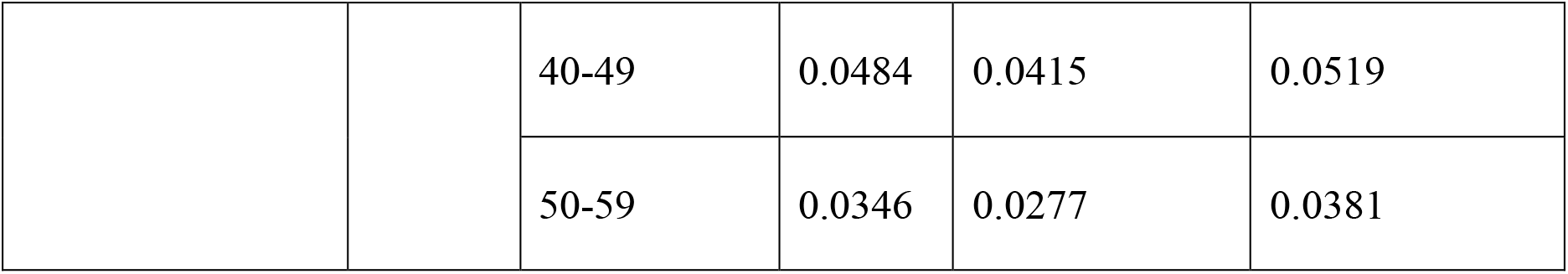
Median values of CSF flow parameters in different conduits by age group.

|  | *Conduit* | *Age Group* | *Median* | *Lower 95% CI* | *Upper 95% CI* |
| --- | --- | --- | --- | --- | --- |
| StroVol (mL) | SA | 20-29 | 0.00217 | 0.00194 | 0.00256 |
|  |  | 30-39 | 0.00288 | 0.00219 | 0.00407 |
|  |  | 40-49 | 0.00303 | 0.00268 | 0.00338 |
|  |  | 50-59 | 0.00190 | 0.00173 | 0.00285 |
|  | COW | 20-29 | 0.0143 | 0.0115 | 0.0157 |
|  |  | 30-39 | 0.0129 | 0.0110 | 0.0143 |
|  |  | 40-49 | 0.0169 | 0.0156 | 0.0192 |
|  |  | 50-59 | 0.0143 | 0.0128 | 0.0157 |
|  | MCA | 20-29 | 0.0121 | 0.0102 | 0.0133 |
|  |  | 30-39 | 0.0176 | 0.0145 | 0.0205 |
|  |  | 40-49 | 0.0174 | 0.0160 | 0.0203 |
|  |  | 50-59 | 0.0118 | 0.0108 | 0.0132 |
| VFR (mL/s) | SA | 20-29 | 0.00600 | 0.00600 | 0.00800 |
|  |  | 30-39 | 0.00700 | 0.00600 | 0.00900 |
|  |  | 40-49 | 0.00700 | 0.00700 | 0.00850 |
|  |  | 50-59 | 0.00600 | 0.00500 | 0.00900 |
|  | COW | 20-29 | 0.0440 | 0.0410 | 0.0480 |
|  |  | 30-39 | 0.0335 | 0.0295 | 0.0370 |
|  |  | 40-49 | 0.0450 | 0.0415 | 0.0480 |
|  |  | 50-59 | 0.0390 | 0.0350 | 0.0450 |
|  | MCA | 20-29 | 0.0375 | 0.0340 | 0.0420 |
|  |  | 30-39 | 0.0545 | 0.0475 | 0.0600 |
|  |  | 40-49 | 0.0530 | 0.0490 | 0.0560 |
|  |  | 50-59 | 0.0370 | 0.0330 | 0.0420 |
| SFR (mL/s) | SA | 20-29 | 0.00800 | 0.00600 | 0.00900 |
|  |  | 30-39 | 0.0100 | 0.00800 | 0.0140 |
|  |  | 40-49 | 0.00900 | 0.00700 | 0.00900 |
|  |  | 50-59 | 0.00800 | 0.00500 | 0.0100 |
|  | COW | 20-29 | 0.0510 | 0.0460 | 0.0560 |
|  |  | 30-39 | 0.0490 | 0.0390 | 0.0580 |
|  |  | 40-49 | 0.0580 | 0.0530 | 0.0640 |
|  |  | 50-59 | 0.0525 | 0.0470 | 0.0580 |
|  | MCA | 20-29 | 0.0500 | 0.0430 | 0.0530 |
|  |  | 30-39 | 0.0620 | 0.0553 | 0.0720 |
|  |  | 40-49 | 0.0660 | 0.0590 | 0.0700 |
|  |  | 50-59 | 0.0470 | 0.0430 | 0.0495 |
| DFR (mL/s) | SA | 20-29 | 0.00350 | 0.00300 | 0.00500 |
|  |  | 30-39 | 0.00400 | 0.00200 | 0.00500 |
|  |  | 40-49 | 0.00500 | 0.00300 | 0.00500 |
|  |  | 50-59 | 0.00400 | 0.00200 | 0.00500 |
|  | COW | 20-29 | 0.0260 | 0.0230 | 0.0330 |
|  |  | 30-39 | 0.0200 | 0.0150 | 0.0230 |
|  |  | 40-49 | 0.0260 | 0.0250 | 0.0330 |
|  |  | 50-59 | 0.0210 | 0.0189 | 0.0240 |
|  | MCA | 20-29 | 0.0270 | 0.0210 | 0.0290 |
|  |  | 30-39 | 0.0285 | 0.0185 | 0.0455 |
|  |  | 40-49 | 0.0310 | 0.0290 | 0.0360 |
|  |  | 50-59 | 0.0190 | 0.0145 | 0.0210 |
| Velocity (cm/s) | SA | 20-29 | 0.556 | 0.455 | 0.578 |
|  |  | 30-39 | 0.540 | 0.442 | 0.589 |
|  |  | 40-49 | 0.674 | 0.578 | 0.783 |
|  |  | 50-59 | 0.723 | 0.578 | 0.867 |
|  | COW | 20-29 | 0.293 | 0.267 | 0.326 |
|  |  | 30-39 | 0.242 | 0.200 | 0.264 |
|  |  | 40-49 | 0.295 | 0.273 | 0.320 |
|  |  | 50-59 | 0.289 | 0.270 | 0.328 |
|  | MCA | 20-29 | 0.267 | 0.248 | 0.289 |
|  |  | 30-39 | 0.273 | 0.248 | 0.318 |
|  |  | 40-49 | 0.269 | 0.253 | 0.293 |
|  |  | 50-59 | 0.237 | 0.217 | 0.255 |
| PSV (cm/s) | SA | 20-29 | 1.16 | 1.16 | 1.73 |
|  |  | 30-39 | 1.30 | 1.30 | 2.31 |
|  |  | 40-49 | 1.45 | 1.45 | 1.88 |
|  |  | 50-59 | 1.59 | 1.16 | 3.47 |
|  | COW | 20-29 | 0.674 | 0.462 | 0.694 |
|  |  | 30-39 | 0.751 | 0.605 | 1.29 |
|  |  | 40-49 | 0.751 | 0.418 | 0.771 |
|  |  | 50-59 | 0.828 | 0.766 | 1.05 |
|  | MCA | 20-29 | 0.535 | 0.484 | 0.815 |
|  |  | 30-39 | 0.676 | 0.124 | 0.700 |
|  |  | 40-49 | 0.766 | 0.489 | 0.867 |
|  |  | 50-59 | 0.605 | 0.500 | 0.867 |
| PDV (cm/s) | SA | 20-29 | 0.867 | 0.247 | 0.867 |
|  |  | 30-39 | 0.578 | 0.578 | 1.76 |
|  |  | 40-49 | 1.07 | 0.867 | 1.54 |
|  |  | 50-59 | 0.723 | 0.578 | 2.75 |
|  | COW | 20-29 | 0.171 | 0.0942 | 0.650 |
|  |  | 30-39 | 0.390 | 0.390 | 0.650 |
|  |  | 40-49 | 0.376 | 0.239 | 0.824 |
|  |  | 50-59 | 0.491 | 0.289 | 0.549 |
|  | MCA | 20-29 | 0.405 | 0.348 | 0.518 |
|  |  | 30-39 | 0.275 | 0.303 | 1.49 |
|  |  | 40-49 | 0.500 | 0.303 | 0.590 |
|  |  | 50-59 | 0.318 | 0.128 | 0.381 |
| TPStroVol (%) | SA | 20-29 | 57.5 | 28.7 | 67.5 |
|  |  | 30-39 | 25.0 | 20.0 | 70.0 |
|  |  | 40-49 | 50.0 | 25.0 | 65.0 |
|  |  | 50-59 | 50.0 | 45.0 | 60.0 |
|  | COW | 20-29 | 75.0 | 70.0 | 92.5 |
|  |  | 30-39 | 70.0 | 70.0 | 90.0 |
|  |  | 40-49 | 70.0 | 67.5 | 90.0 |
|  |  | 50-59 | 75.0 | 50.0 | 75.0 |
|  | MCA | 20-29 | 65.0 | 65.0 | 80.0 |
|  |  | 30-39 | 65.0 | 65.0 | 90.0 |
|  |  | 40-49 | 75.0 | 50.0 | 75.0 |
|  |  | 50-59 | 70.0 | 70.0 | 90.0 |
| TPVFR (%) | SA | 20-29 | 17.5 | 10.0 | 35.0 |
|  |  | 30-39 | 22.5 | 20.0 | 67.5 |
|  |  | 40-49 | 22.5 | 12.5 | 25.0 |
|  |  | 50-59 | 20.0 | 5.00 | 55.0 |
|  | COW | 20-29 | 50.0 | 27.5 | 60.0 |
|  |  | 30-39 | 42.5 | 32.5 | 70.0 |
|  |  | 40-49 | 25.0 | 20.0 | 55.0 |
|  |  | 50-59 | 50.0 | 20.0 | 60.0 |
|  | MCA | 20-29 | 47.5 | 35.0 | 65.0 |
|  |  | 30-39 | 45.0 | 32.5 | 65.0 |
|  |  | 40-49 | 15.0 | 15.0 | 60.0 |
|  |  | 50-59 | 35.0 | 15.0 | 45.0 |
| TPVelocity (%) | SA | 20-29 | 55.0 | 50.0 | 72.5 |
|  |  | 30-39 | 50.0 | 50.0 | 90.0 |
|  |  | 40-49 | 55.0 | 55.0 | 85.0 |
|  |  | 50-59 | 50.0 | 40.0 | 55.0 |
|  | COW | 20-29 | 60.0 | 35.0 | 75.0 |
|  |  | 30-39 | 80.0 | 60.0 | 90.0 |
|  |  | 40-49 | 57.5 | 50.0 | 72.5 |
|  |  | 50-59 | 60.0 | 45.0 | 80.0 |
|  | MCA | 20-29 | 60.0 | 45.0 | 90.0 |
|  |  | 30-39 | 57.5 | 30.0 | 75.0 |
|  |  | 40-49 | 75.0 | 45.0 | 80.0 |
|  |  | 50-59 | 70.0 | 45.0 | 80.0 |
| Area (cm<sup>2</sup>) | SA | 20-29 | 0.0104 | 0.00692 | 0.0104 |
|  |  | 30-39 | 0.0121 | 0.0104 | 0.0173 |
|  |  | 40-49 | 0.0104 | 0.00692 | 0.0104 |
|  |  | 50-59 | 0.00692 | 0.00346 | 0.00692 |
|  | COW | 20-29 | 0.0311 | 0.0242 | 0.0346 |
|  |  | 30-39 | 0.0311 | 0.0260 | 0.0415 |
|  |  | 40-49 | 0.0311 | 0.0277 | 0.0381 |
|  |  | 50-59 | 0.0208 | 0.0208 | 0.0293 |
|  | MCA | 20-29 | 0.0311 | 0.0294 | 0.0381 |
|  |  | 30-39 | 0.0588 | 0.0484 | 0.0813 |
|  |  | 40-49 | 0.0484 | 0.0415 | 0.0519 |
|  |  | 50-59 | 0.0346 | 0.0277 | 0.0381 |

**Table 5.** Estimates of models of CSF flow parameters in SA (intercept) vs. COW vs. MCA.

|  | *Fixed Effects* | *Coefficient* | *Lower*<br>*95% CI* | *Upper*<br>*95% CI* | *Random*<br>*Effects* | *Variance* | *SD* | *H0* | *P-Value* |
| --- | --- | --- | --- | --- | --- | --- | --- | --- | --- |
| StroVol | Intercept (SA) | -0.00857 | -0.0124 | -0.00527 | Intercept:<br>Participants | 5.22E-06 | 0.00228 | SA = COW = MCA | 0.000999 |
|  | Time | 0.0278 | 0.0207 | 0.0335 | Residuals | 0.00142 | 0.0376 | SA = COW | 0.00400 |
|  | COW | 0.0150 | 0.0103 | 0.0190 |  |  |  | SA = MCA | 0.00400 |
|  | MCA | 0.0142 | 0.00979 | 0.0194 |  |  |  | COW = MCA | 0.805 |
| VFR | Intercept (SA) | 0.0230 | 0.0164 | 0.0303 | Intercept:<br>Participants | 2.24E-05 | 0.00474 | SA = COW = MCA | 0.000999 |
|  | Time | -0.0241 | -0.0363 | -0.0139 | Residuals | 0.00598 | 0.0773 | SA = COW | 0.00400 |
|  | COW | 0.0350 | 0.0261 | 0.0460 |  |  |  | SA = MCA | 0.00400 |
|  | MCA | 0.0310 | 0.0203 | 0.0411 |  |  |  | COW = MCA | 0.529 |
| SFR | Intercept (SA) | -3.73 | -3.88 | -3.63 | Intercept:<br>Participants | 0.0172 | 0.131 | SA = COW = MCA | 0.00100 |
|  | Time | -0.483 | -0.705 | -0.320 | Residuals | 1.07 | 1.03 | SA = COW | 0.00400 |
|  | COW | 1.36 | 1.20 | 1.50 |  |  |  | SA = MCA | 0.00400 |
|  | MCA | 1.36 | 1.21 | 1.51 |  |  |  | COW = MCA | 0.969 |
| DFR | Intercept (SA) | -4.78 | -5.17 | -4.55 | Intercept:<br>Participants | 0.0738 | 0.272 | SA = COW = MCA | 0.00100 |
|  | Time | -0.536 | -0.802 | -0.245 | Residuals | 0.816 | 0.903 | SA = COW | 0.00799 |
|  | COW | 1.48 | 1.16 | 1.83 |  |  |  | SA = MCA | 0.00799 |
|  | MCA | 1.61 | 1.33 | 1.94 |  |  |  | COW = MCA | 0.216 |
| Velocity | Intercept (SA) | 0.421 | 0.318 | 0.534 | Intercept:<br>Participants | 0.00180 | 0.0424 | SA = COW = MCA | 0.000999 |
|  | Time | -0.104 | -0.178 | -0.0277 | Residuals | 0.308 | 0.555 | SA = COW | 0.0160 |
|  | COW | -0.159 | -0.286 | -0.0498 |  |  |  | SA = MCA | 0.00599 |
|  | MCA | -0.198 | 0.0204 | 0.0410 |  |  |  | COW = MCA | 0.166 |
| PSV | Intercept (SA) | 0.537 | 0.413 | 0.645 |  |  |  | SA = COW = MCA | 0.000999 |
|  | COW | -0.773 | -0.954 | -0.613 |  |  |  | SA = COW | 0.00400 |
|  | MCA | -0.955 | -1.12 | -0.773 |  |  |  | SA = MCA | 0.00400 |
|  |  |  |  |  |  |  |  | COW = MCA | 0.0280 |
| PDV | Intercept (SA) | 0.184 | -0.0516 | 0.378 |  |  |  | SA = COW = MCA | 0.000999 |
|  | COW | -0.984 | -1.24 | -0.721 |  |  |  | SA = COW | 0.00400 |
|  | MCA | -1.01 | -1.29 | -0.747 |  |  |  | SA = MCA | 0.00400 |
|  |  |  |  |  |  |  |  | COW = MCA | 0.901 |
| TPStroVol | Intercept (SA)<br>(Mean Model) | -0.0633 | -0.329 | 0.202 |  |  |  | SA = COW = MCA | 3.87E-10 |
|  | COW (Mean Model) | 1.12 | 0.792 | 1.45 |  |  |  | SA = COW | 7.34E-11 |
|  | MCA (Mean Model) | 0.966 | 0.646 | 1.29 |  |  |  | SA = MCA | 1.19E-08 |
|  | Intercept (SA)<br>(Precision Model) | 1.21 | 0.859 | 1.57 |  |  |  | COW = MCA | 0.482 |
|  | COW (Precision Model) | 0.590 | 0.125 | 1.06 |  |  |  |  |  |
|  | MCA (Precision Model) | 0.746 | 0.274 | 1.22 |  |  |  |  |  |
| TPVFR | Intercept (SA)<br>(Mean Model) | -0.979 | -1.26 | -0.696 |  |  |  | SA = COW = MCA | 0.00131 |
|  | COW (Mean Model) | 0.690 | 0.322 | 1.06 |  |  |  | SA = COW | 0.000468 |
|  | MCA (Mean Model) | 0.486 | 0.110 | 0.862 |  |  |  | SA = MCA | 0.0274 |
|  | Intercept (SA)<br>(Precision Model) | 1.33 | 0.959 | 1.70 |  |  |  | COW = MCA | 0.471 |
|  | COW (Precision Model) | -0.450 | -0.910 | 0.0100 |  |  |  |  |  |
|  | MCA (Precision Model) | -0.434 | -0.902 | 0.0342 |  |  |  |  |  |
| TPVelocity | Intercept (SA)<br>(Mean Model) | 0.465 | 0.288 | 0.642 |  |  |  | SA = COW = MCA | 0.861 |
|  | COW (Mean Model) | 0.0137 | -0.286 | 0.313 |  |  |  |  |  |
|  | MCA (Mean Model) | 0.0846 | -0.224 | 0.393 |  |  |  |  |  |
|  | Intercept (SA) (Precision Model) | 1.60 | 1.32 | 1.88 |  |  |  |  |  |
|  | COW (Precision Model) | -0.820 | -1.20 | -0.436 |  |  |  |  |  |
|  | MCA (Precision Model) | -0.762 | -1.16 | -0.366 |  |  |  |  |  |
| Area | Intercept (SA) | -2.81 | -2.96 | -2.66 | Intercept: Participants | 0.135 | 0.367 | SA = COW = MCA | 0.000999 |
|  | Time | -1.43 | -1.67 | -1.22 | Residuals | 2.89 | 1.70 | SA = COW | 0.00400 |
|  | COW | 0.984 | 0.793 | 1.15 |  |  |  | SA = MCA | 0.00400 |
|  | MCA | 1.08 | 0.867 | 1.27 |  |  |  | COW = MCA | 0.230 |

**Table 6.** Estimates of models of CSF flow parameters vs. sex, age, and their interaction for each conduit.

|  | *Conduit* | *Fixed Effects* | *Coefficient* | *Lower 95% CI* | *Upper 95% CI* | *Random Effects* | *Variance* | *SD* | *H0* | *P-Value* |
| --- | --- | --- | --- | --- | --- | --- | --- | --- | --- | --- |
| StroVol | SA | Intercept | 0.00554 | 0.000963 | 0.0103 | Intercept: Participants | 2.37E-06 | 0.00154 | Sex = 0, Age = 0, and Sex:Age = 0 | 0.694 |
|  |  | Time | -0.00133 | -0.00385 | 0.00152 | Residuals | 9.57E-05 | 0.00978 |  |  |
|  |  | Sex | -0.00164 | -0.00718 | 0.00459 |  |  |  |  |  |
|  |  | Age | -2.57E-05 | -0.000152 | 9.55E-05 |  |  |  |  |  |
|  |  | Sex x Age | 1.68E-05 | -0.000141 | 0.000154 |  |  |  |  |  |
|  | COW | Intercept | 0.0125 | -0.0151 | 0.0369 | Intercept: Participants | 7.28E-06 | 0.00270 | Sex = 0, Age = 0, and Sex:Age = 0 | 0.385 |
|  |  | Time | 0.0309 | 0.0206 | 0.0399 | Residuals | 0.00154 | 0.0392 |  |  |
|  |  | Sex | -0.00549 | -0.0343 | 0.0318 |  |  |  |  |  |
|  |  | Age | -0.000201 | -0.000828 | 0.000445 |  |  |  |  |  |
|  |  | Sex x Age | 0.000164 | -0.000551 | 0.000933 |  |  |  |  |  |
|  | MCA | Intercept | -0.00496 | -0.0262 | 0.0192 | Intercept:<br>Participants | 2.69E-05 | 0.00519 | Sex = 0, Age = 0,<br>and Sex:Age = 0 | 0.334 |
|  |  | Time | 0.0345 | 0.0239 | 0.0455 | Residuals | 0.00168 | 0.0410 |  |  |
|  |  | Sex | 0.0177 | -0.0128 | 0.0488 |  |  |  |  |  |
|  |  | Age | 0.000171 | -0.000441 | 0.000683 |  |  |  |  |  |
|  |  | Sex x Age | -0.000393 | -0.00115 | 0.000376 |  |  |  |  |  |
| VFR | SA | Intercept | 0.0244 | 0.00850 | 0.0383 | Intercept:<br>Participants | 1.57E-05 | 0.00397 | Sex = 0, Age = 0,<br>and Sex:Age = 0 | 0.835 |
|  |  | Time | -0.0234 | -0.0334 | -0.0151 | Residuals | 0.000719 | 0.0268 |  |  |
|  |  | Sex | -0.00787 | -0.0265 | 0.0104 |  |  |  |  |  |
|  |  | Age | -4.93E-05 | -0.000414 | 0.000367 |  |  |  |  |  |
|  |  | Sex x Age | 0.000185 | -0.000298 | 0.000661 |  |  |  |  |  |
|  | COW | Intercept | 0.0757 | 0.0178 | 0.133 | Intercept:<br>Participants | 4.00E-05 | 0.00632 | Sex = 0, Age = 0,<br>and Sex:Age = 0 | 0.445 |
|  |  | Time | -0.0301 | -0.0479 | -0.0148 | Residuals | 0.00653 | 0.0808 |  |  |
|  |  | Sex | -0.0233 | -0.101 | 0.0538 |  |  |  |  |  |
|  |  | Age | -0.000384 | -0.00194 | 0.00107 |  |  |  |  |  |
|  |  | Sex x Age | 0.000590 | -0.00117 | 0.00243 |  |  |  |  |  |
|  | MCA | Intercept | 0.0257 | -0.0223 | 0.0812 | Intercept:<br>Participants | 0.000101 | 0.0101 | Sex = 0, Age = 0,<br>and Sex:Age = 0 | 0.0879 |
|  |  | Time | -0.0179 | -0.0382 | 0.00150 | Residuals | 0.00705 | 0.0840 |  |  |
|  |  | Sex | 0.0507 | -0.0138 | 0.122 |  |  |  |  |  |
|  |  | Age | 0.000574 | -0.000657 | 0.00178 |  |  |  |  |  |
|  |  | Sex x Age | -0.00112 | -0.00283 | 0.000421 |  |  |  |  |  |
| SFR | SA | Intercept | -3.24 | -3.52 | -3.00 | Intercept:<br>Participants | 0.0156 | 0.125 | Sex = 0, Age = 0,<br>and Sex:Age = 0 | 0.787 |
|  |  | Time | -1.61 | -2.35 | -1.18 | Residuals | 1.93 | 1.39 |  |  |
|  |  | Sex | -0.207 | -0.447 | 0.0393 |  |  |  |  |  |
|  |  | Age | -0.0190 | -0.222 | 0.139 |  |  |  |  |  |
|  |  | Sex x Age | 0.0246 | -0.234 | 0.299 |  |  |  |  |  |
|  | COW | Intercept | -2.38 | -2.62 | -2.28 | Intercept:<br>Participants | 0.0135 | 0.116 | Sex = 0, Age = 0,<br>and Sex:Age = 0 | 0.0539 |
|  |  | Time | -0.368 | -0.552 | -0.175 | Residuals | 0.967 | 0.983 |  |  |
|  |  | Sex | -0.104 | -0.370 | 0.105 |  |  |  |  |  |
|  |  | Age | -0.0128 | -0.179 | 0.169 |  |  |  |  |  |
|  |  | Sex x Age | 0.0625 | -0.155 | 0.305 |  |  |  |  |  |
|  | MCA | Intercept | -2.43 | -2.67 | -2.34 | Intercept:<br>Participants | 0.0466 | 0.216 | Sex = 0, Age = 0,<br>and Sex:Age = 0 | 0.157 |
|  |  | Time | -0.329 | -0.581 | -0.110 | Residuals | 0.843 | 0.918 |  |  |
|  |  | Sex | -0.0412 | -0.262 | 0.187 |  |  |  |  |  |
|  |  | Age | 0.0355 | -0.186 | 0.215 |  |  |  |  |  |
|  |  | Sex x Age | -0.150 | -0.386 | 0.0833 |  |  |  |  |  |
| DFR | SA | Intercept | -4.67 | -4.67 | -4.04 | Intercept:<br>Participants | 0.140 | 0.374 | Sex = 0, Age = 0,<br>and Sex:Age = 0 | 0.996 |
|  |  | Time | -1.32 | -1.61 | -0.761 | Residuals | 0.783 | 0.885 |  |  |
|  |  | Sex | 0.433 | -0.837 | -0.194 |  |  |  |  |  |
|  |  | Age | 0.0105 | -0.0999 | 0.394 |  |  |  |  |  |
|  |  | Sex x Age | -0.0251 | -0.587 | 0.0329 |  |  |  |  |  |
|  | COW | Intercept | -3.32 | -3.63 | -3.15 | Intercept:<br>Participants | 0.0315 | 0.177 | Sex = 0, Age = 0,<br>and Sex:Age = 0 | 0.548 |
|  |  | Time | -0.253 | -0.588 | 0.0993 | Residuals | 0.661 | 0.813 |  |  |
|  |  | Sex | -0.266 | -0.435 | -0.0693 |  |  |  |  |  |
|  |  | Age | -0.00338 | -0.161 | 0.174 |  |  |  |  |  |
|  |  | Sex x Age | -0.0578 | -0.197 | 0.145 |  |  |  |  |  |
|  | MCA | Intercept | -3.22 | -3.48 | -3.07 | Intercept:<br>Participants | 0.100 | 0.317 | Sex = 0, Age = 0,<br>and Sex:Age = 0 | 0.999 |
|  |  | Time | -0.293 | -0.636 | 0.0317 | Residuals | 0.745 | 0.863 |  |  |
|  |  | Sex | -0.136 | -0.425 | 0.156 |  |  |  |  |  |
|  |  | Age | 0.0760 | -0.194 | 0.289 |  |  |  |  |  |
|  |  | Sex x Age | -0.287 | -0.541 | -0.00865 |  |  |  |  |  |
| Velocity | SA | Intercept | 0.0460 | -0.851 | 1.13 | Intercept:<br>Participants | 0.0128 | 0.113 | Sex = 0, Age = 0,<br>and Sex:Age = 0 | 0.217 |
|  |  | Time | -0.729 | -1.03 | -0.409 | Residuals | 1.34 | 1.16 |  |  |
|  |  | Sex | 0.469 | -0.640 | 1.32 |  |  |  |  |  |
|  |  | Age | 0.0182 | -0.0113 | 0.0410 |  |  |  |  |  |
|  |  | Sex x Age | -0.0156 | -0.0431 | 0.0150 |  |  |  |  |  |
|  | COW | Intercept | 0.352 | 0.0471 | 0.684 | Intercept:<br>Participants | 0.00236 | 0.0486 | Sex = 0, Age = 0,<br>and Sex:Age = 0 | 0.249 |
|  |  | Time | -0.00946 | -0.0934 | 0.0795 | Residuals | 0.146 | 0.382 |  |  |
|  |  | Sex | -0.0995 | -0.481 | 0.306 |  |  |  |  |  |
|  |  | Age | -0.00347 | -0.0114 | 0.00469 |  |  |  |  |  |
|  |  | Sex x Age | 0.00285 | -0.00672 | 0.0120 |  |  |  |  |  |
|  | MCA | Intercept | 0.0482 | -0.202 | 0.280 | Intercept:<br>Participants | 0.00372 | 0.0610 | Sex = 0, Age = 0,<br>and Sex:Age = 0 | 0.273 |
|  |  | Time | 0.0113 | -0.0777 | 0.0977 | Residuals | 0.112 | 0.335 |  |  |
|  |  | Sex | 0.241 | -0.0499 | 0.555 |  |  |  |  |  |
|  |  | Age | 0.00292 | -0.00226 | 0.00896 |  |  |  |  |  |
|  |  | Sex x Age | -0.00554 | -0.0123 | 0.00113 |  |  |  |  |  |
| PSV | SA | Intercept | 0.540 | -0.356 | 1.41 |  |  |  | Sex = 0, Age = 0,<br>and Sex:Age = 0 | 0.0579 |
|  |  | Sex | -0.520 | -1.54 | 0.457 |  |  |  |  |  |
|  |  | Age | 0.00399 | -0.0195 | 0.0242 |  |  |  |  |  |
|  |  | Sex x Age | 0.00177 | -0.0233 | 0.0248 |  |  |  |  |  |
|  | COW | Intercept | 0.0723 | -0.623 | 0.647 |  |  |  | Sex = 0, Age = 0,<br>and Sex:Age = 0 | 0.273 |
|  |  | Sex | -0.838 | -1.62 | 0.0779 |  |  |  |  |  |
|  |  | Age | -0.00625 | -0.0230 | 0.0102 |  |  |  |  |  |
|  |  | Sex x Age | 0.0180 | -0.00132 | 0.0387 |  |  |  |  |  |
|  | MCA | Intercept | -0.528 | -1.16 | 0.126 |  |  |  | Sex = 0, Age = 0,<br>and Sex:Age = 0 | 0.208 |
|  |  | Sex | -0.141 | -1.16 | 0.666 |  |  |  |  |  |
|  |  | Age | 0.00504 | -0.0121 | 0.0206 |  |  |  |  |  |
|  |  | Sex x Age | -0.00195 | -0.0207 | 0.0218 |  |  |  |  |  |
| PDV | SA | Intercept | -0.133 | -2.18 | 1.11 |  |  |  | Sex = 0, Age = 0,<br>and Sex:Age = 0 | 0.000999 |
|  |  | Sex | -0.0369 | -1.41 | 2.17 |  |  |  | Sex = 0 | 0.308 |
|  |  | Age | 0.0167 | -0.0179 | 0.0607 |  |  |  | Age = 0 | 0.308 |
|  |  | Sex x Age | -0.0199 | -0.0754 | 0.0163 |  |  |  | Sex:Age = 0 | 0.308 |
|  | COW | Intercept | -0.820 | -2.73 | 0.227 |  |  |  | Sex = 0, Age = 0,<br>and Sex:Age = 0 | 0.0180 |
|  |  | Sex | -1.12 | -2.72 | 0.644 |  |  |  | Sex = 0 | 0.647 |
|  |  | Age | 0.00387 | -0.0236 | 0.0447 |  |  |  | Age = 0 | 0.827 |
|  |  | Sex x Age | 0.0197 | -0.0243 | 0.0573 |  |  |  | Sex:Age = 0 | 0.647 |
|  | MCA | Intercept | -0.240 | -0.986 | 0.833 |  |  |  | Sex = 0, Age = 0,<br>and Sex:Age = 0 | 0.595 |
|  |  | Sex | -0.841 | -2.28 | 0.546 |  |  |  |  |  |
|  |  | Age | -0.0133 | -0.0450 | 0.00558 |  |  |  |  |  |
|  |  | Sex x Age | 0.0179 | -0.0182 | 0.0492 |  |  |  |  |  |
| TPStroVol | SA | Intercept<br>(Mean<br>Model) | 0.932 | -0.651 | 2.52 |  |  |  | Sex = 0, Age = 0,<br>and Sex:Age = 0 | 0.164 |
|  |  | Sex<br>(Mean<br>Model) | -1.25 | -3.27 | 0.777 |  |  |  |  |  |
|  |  | Age<br>(Mean<br>Model) | -0.0183 | -0.0572 | 0.0207 |  |  |  |  |  |
|  |  | Sex x Age<br>(Mean<br>Model) | 0.0179 | -0.0338 | 0.0697 |  |  |  |  |  |
|  |  | Intercept<br>(Precision<br>Model) | 0.140 | -1.79 | 2.07 |  |  |  |  |  |
|  |  | Sex<br>(Precision<br>Model) | 2.67 | -0.154 | 5.50 |  |  |  |  |  |
|  |  | Age<br>(Precision<br>Model) | 0.0267 | -0.0232 | 0.0765 |  |  |  |  |  |
|  |  | Sex x Age<br>(Precision<br>Model) | -0.0554 | -0.127 | 0.0160 |  |  |  |  |  |
|  | COW | Intercept<br>(Mean<br>Model) | 1.91 | 0.599 | 3.23 |  |  |  | Sex = 0, Age = 0,<br>and Sex:Age = 0 | 0.708 |
|  |  | Sex<br>(Mean<br>Model) | -0.600 | -2.33 | 1.13 |  |  |  |  |  |
|  |  | Age<br>(Mean<br>Model) | -0.0181 | -0.0490 | 0.0128 |  |  |  |  |  |
|  |  | Sex x Age<br>(Mean<br>Model) | 0.0150 | -0.0265 | 0.0566 |  |  |  |  |  |
|  |  | Intercept<br>(Precision<br>Model) | 1.62 | -0.343 | 3.59 |  |  |  |  |  |
|  |  | Sex<br>(Precision<br>Model) | 0.377 | -2.14 | 2.90 |  |  |  |  |  |
|  |  | Age<br>(Precision<br>Model) | 0.00488 | -0.0423 | 0.0520 |  |  |  |  |  |
|  |  | Sex x Age<br>(Precision Model) | -0.0158 | -0.0764 | 0.0448 |  |  |  |  |  |
|  | MCA | Intercept<br>(Mean Model) | 1.69 | 0.582 | 2.80 |  |  |  | Sex = 0, Age = 0,<br>and Sex:Age = 0 | 0.0282 |
|  |  | Sex<br>(Mean Model) | -1.50 | -2.87 | -0.126 |  |  |  | Sex = 0 | 0.108 |
|  |  | Age<br>(Mean Model) | -0.0127 | -0.0405 | 0.0150 |  |  |  | Age = 0 | 0.374 |
|  |  | Sex x Age<br>(Mean Model) | 0.0284 | -0.00728 | 0.0640 |  |  |  | Sex:Age = 0 | 0.158 |
|  |  | Intercept<br>(Precision Model) | 3.16 | 1.20 | 5.12 |  |  |  |  |  |
|  |  | Sex<br>(Precision Model) | 0.874 | -1.69 | 3.44 |  |  |  |  |  |
|  |  | Age<br>(Precision Model) | -0.0305 | -0.0775 | 0.0165 |  |  |  |  |  |
|  |  | Sex x Age<br>(Precision Model) | -0.0158 | -0.0769 | 0.0453 |  |  |  |  |  |
| TPVFR | SA | Intercept<br>(Mean Model) | -0.666 | -2.25 | 0.915 |  |  |  | Sex = 0, Age = 0,<br>and Sex:Age = 0 | 0.913 |
|  |  | Sex<br>(Mean<br>Model) | 0.0741 | -2.14 | 2.29 |  |  |  |  |  |
|  |  | Age<br>(Mean<br>Model) | -0.00940 | -0.0504 | 0.0316 |  |  |  |  |  |
|  |  | Sex x Age<br>(Mean<br>Model) | 0.000817 | -0.0568 | 0.0584 |  |  |  |  |  |
|  |  | Intercept<br>(Precision<br>Model) | 1.38 | -0.687 | 3.45 |  |  |  |  |  |
|  |  | Sex<br>(Precision<br>Model) | 0.373 | -2.56 | 3.31 |  |  |  |  |  |
|  |  | Age<br>(Precision<br>Model) | -0.00181 | -0.0550 | 0.0514 |  |  |  |  |  |
|  |  | Sex x Age<br>(Precision<br>Model) | -0.00762 | -0.0826 | 0.0673 |  |  |  |  |  |
|  | COW | Intercept<br>(Mean<br>Model) | 0.0577 | -1.24 | 1.36 |  |  |  | Sex = 0, Age = 0,<br>and Sex:Age = 0 | 0.0912 |
|  |  | Sex<br>(Mean<br>Model) | 0.233 | -1.54 | 2.01 |  |  |  |  |  |
|  |  | Age<br>(Mean<br>Model) | -0.0161 | -0.0493 | 0.0171 |  |  |  |  |  |
|  |  | Sex x Age<br>(Mean Model) | 0.00874 | -0.0351 | 0.0526 |  |  |  |  |  |
|  |  | Intercept<br>(Precision Model) | 2.48 | 0.662 | 4.31 |  |  |  |  |  |
|  |  | Sex<br>(Precision Model) | -1.95 | -4.25 | 0.353 |  |  |  |  |  |
|  |  | Age<br>(Precision Model) | -0.0345 | -0.0782 | 0.00921 |  |  |  |  |  |
|  |  | Sex x Age<br>(Precision Model) | 0.0439 | -0.0116 | 0.0995 |  |  |  |  |  |
|  | MCA | Intercept<br>(Mean Model) | 0.339 | -1.14 | 1.81 |  |  |  | Sex = 0, Age = 0,<br>and Sex:Age = 0 | 0.587 |
|  |  | Sex<br>(Mean Model) | -0.780 | -2.72 | 1.16 |  |  |  |  |  |
|  |  | Age<br>(Mean Model) | -0.0227 | -0.0592 | 0.0137 |  |  |  |  |  |
|  |  | Sex x Age<br>(Mean Model) | 0.0240 | -0.0237 | 0.0717 |  |  |  |  |  |
|  |  | Intercept<br>(Precision Model) | 1.33 | -0.439 | 3.11 |  |  |  |  |  |
|  |  | Sex<br>(Precision<br>Model) | 0.157 | -2.17 | 2.49 |  |  |  |  |  |
|  |  | Age<br>(Precision<br>Model) | -0.0101 | -0.0530 | 0.0328 |  |  |  |  |  |
|  |  | Sex x Age<br>(Precision<br>Model) | -0.00328 | -0.0591 | 0.0526 |  |  |  |  |  |
| TPVelocity | SA | Intercept<br>(Mean<br>Model) | 0.488 | -0.753 | 1.73 |  |  |  | Sex = 0, Age = 0,<br>and Sex:Age = 0 | 0.115 |
|  |  | Sex<br>(Mean<br>Model) | -0.801 | -2.32 | 0.720 |  |  |  |  |  |
|  |  | Age<br>(Mean<br>Model) | 0.00229 | -0.0276 | 0.0322 |  |  |  |  |  |
|  |  | Sex x Age<br>(Mean<br>Model) | 0.0154 | -0.0249 | 0.0556 |  |  |  |  |  |
|  |  | Intercept<br>(Precision<br>Model) | -0.283 | -2.01 | 1.44 |  |  |  |  |  |
|  |  | Sex<br>(Precision<br>Model) | 4.60 | 2.32 | 6.88 |  |  |  |  |  |
|  |  | Age<br>(Precision<br>Model) | 0.0492 | 0.00490 | 0.0934 |  |  |  |  |  |
|  |  | Sex x Age<br>(Precision<br>Model) | -0.112 | -0.171 | -0.0541 |  |  |  |  |  |
|  | COW | Intercept<br>(Mean<br>Model) | -0.0387 | -1.53 | 1.46 |  |  |  | Sex = 0, Age = 0,<br>and Sex:Age = 0 | 0.860 |
|  |  | Sex<br>(Mean<br>Model) | 0.868 | -1.10 | 2.83 |  |  |  |  |  |
|  |  | Age<br>(Mean<br>Model) | 0.0119 | -0.0230 | 0.0468 |  |  |  |  |  |
|  |  | Sex x Age<br>(Mean<br>Model) | -0.0198 | -0.0665 | 0.0269 |  |  |  |  |  |
|  |  | Intercept<br>(Precision<br>Model) | -0.190 | -1.93 | 1.55 |  |  |  |  |  |
|  |  | Sex<br>(Precision<br>Model) | 1.07 | -1.11 | 3.26 |  |  |  |  |  |
|  |  | Age<br>(Precision<br>Model) | 0.0321 | -0.0104 | 0.0746 |  |  |  |  |  |
|  |  | Sex x Age<br>(Precision<br>Model) | -0.0392 | -0.0923 | 0.0138 |  |  |  |  |  |
|  | MCA | Intercept<br>(Mean<br>Model) | 0.989 | -0.586 | 2.56 |  |  |  | Sex = 0, Age = 0,<br>and Sex:Age = 0 | 0.827 |
|  |  | Sex<br>(Mean<br>Model) | -0.562 | -2.61 | 1.49 |  |  |  |  |  |
|  |  | Age<br>(Mean<br>Model) | -0.00819 | -0.0457 | 0.0294 |  |  |  |  |  |
|  |  | Sex x Age<br>(Mean<br>Model) | 0.00846 | -0.0409 | 0.0579 |  |  |  |  |  |
|  |  | Intercept<br>(Precision<br>Model) | 0.713 | -1.09 | 2.52 |  |  |  |  |  |
|  |  | Sex<br>(Precision<br>Model) | 0.509 | -1.80 | 2.82 |  |  |  |  |  |
|  |  | Age<br>(Precision<br>Model) | 0.00586 | -0.0375 | 0.0493 |  |  |  |  |  |
|  |  | Sex x Age<br>(Precision<br>Model) | -0.0174 | -0.0729 | 0.0381 |  |  |  |  |  |
| Area | SA | Intercept | -1.19 | -2.38 | -0.604 | Intercept:<br>Participants | 0.392 | 0.626 | Sex = 0, Age = 0,<br>and Sex:Age = 0 | 1.00 |
|  |  | Time | -2.76 | -3.22 | -2.38 | Residuals | 2.77 | 1.66 |  |  |
|  |  | Sex | -0.752 | -1.56 | 0.423 |  |  |  |  |  |
|  |  | Age | -0.0337 | -0.0498 | -0.00877 |  |  |  |  |  |
|  |  | Sex x Age | 0.0231 | -0.0112 | 0.0459 |  |  |  |  |  |
|  | COW | Intercept | -1.86 | -2.13 | -1.69 | Intercept: Participants | 0.262 | 0.512 | Sex = 0, Age = 0, and Sex:Age = 0 | 1.00 |
|  |  | Time | -1.50 | -1.93 | -1.21 | Residuals | 3.16 | 1.78 |  |  |
|  |  | Sex | 0.0753 | -0.160 | 0.311 |  |  |  |  |  |
|  |  | Age | 0.0380 | -0.133 | 0.189 |  |  |  |  |  |
|  |  | Sex x Age | -0.120 | -0.361 | 0.0939 |  |  |  |  |  |
|  | MCA | Intercept | -2.35 | -2.97 | -1.76 | Intercept: Participants | 0.283 | 0.532 | Sex = 0, Age = 0, and Sex:Age = 0 | 1.00 |
|  |  | Time | -0.941 | -1.30 | -0.681 | Residuals | 2.46 | 1.57 |  |  |
|  |  | Sex | 0.954 | -0.0166 | 1.82 |  |  |  |  |  |
|  |  | Age | 0.00874 | -0.00643 | 0.0242 |  |  |  |  |  |
|  |  | Sex x Age | -0.0231 | -0.0430 | -0.000284 |  |  |  |  |  |

As expected, VFR oscillates in the SA which reflects cardiac pulsations. On the other hand, COW VFR decreases throughout the cardiac cycle (Fig. 2a). Interestingly, Velocity in the SA is pulsatile while it stays relatively constant in the COW and MCA (Fig. 2b).

**Figure 2.**
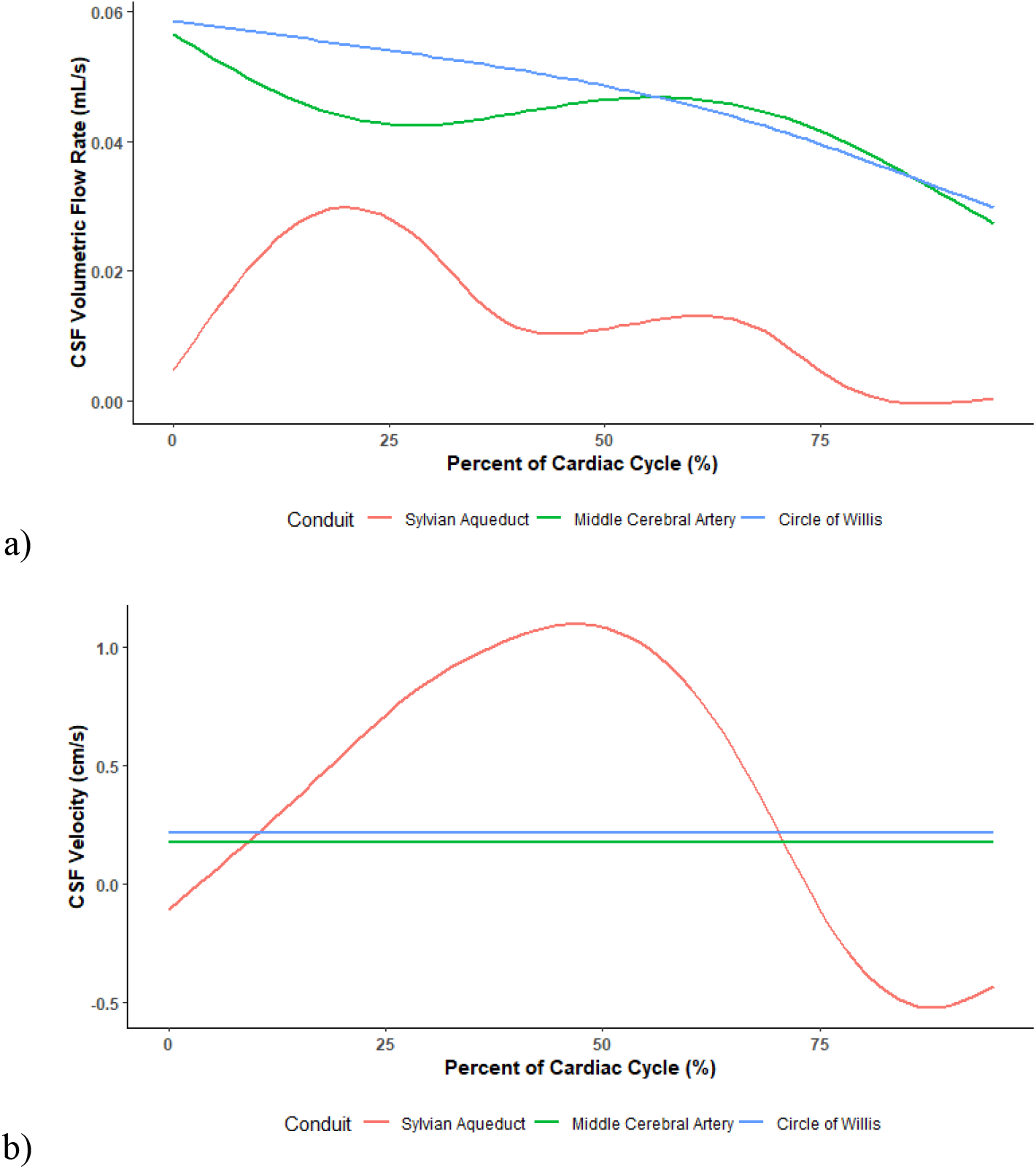
Aggregate CSF flow curves. Aggregation of CSF (a) VFR and (b) Velocity curves for all participants.

SA was set as the intercept because CSF flow parameters for this conduit are already established in the literature. The StroVol LMM indicates strong effects with COW StroVol 0.0150 mL (95% CI [0.0103, 0.0190], *p* = 0.00400) and MCA StroVol 0.0142 mL (95% CI [0.00979, 0.0194], *p* = 0.00400) higher than SA (Table 5). The TPStroVol occurs later in the cardiac cycle in COW (β = 1.12, 95% CI [0.792, 1.45], *p* = 7.34 x 10^-11^) and MCA (β = 0.966, 95% CI [0.646, 1.29], *p* = 1.19 x 10^-8^) compared to SA (Table 5).

The LMMs and GLMMs for flow also indicate strong effects of the conduit as the magnitude of the parameters is higher in COW and MCA. Specifically, COW VFR is 0.0350 mL/s (95% CI [0.0261, 0.0460], *p* = 0.00400) and MCA VFR is 0.0310 mL/s (95% CI [0.0203, 0.0411], *p* = 0.00400) higher than SA (Table 5). COW SFR increases by a factor of 1.36 (95% CI [1.20, 1.50], *p* = 0.00400) and MCA SFR by a factor of 1.36 (95% CI [1.21, 1.51], *p* = 0.00400) compared to SA (Table 5). COW DFR also increases by a factor of 1.48 (95% CI [1.16, 1.83], *p* = 0.00799) and MCA SFR by a factor of 1.61 (95% CI [1.33, 1.94], *p* = 0.00799) compared to SA (Table 5). The TPVFR also occurs later in COW (β = 0.690, 95% CI [0.322, 1.06], *p* = 0.000468) and MCA (β = 0.486, 95% CI [0.110, 0.862], *p* = 0.0274) compared to SA (Table 5).

Velocity in the LMM and GLMs also exhibits strong effects, and the magnitude of the parameters is lower in COW and MCA. COW Velocity is 0.159 cm/s (95% CI [-0.286, −0.0498], *p* = 0.0160) and MCA Velocity is 0.198 cm/s (95% CI [0.0204, 0.0410], *p* = 0.00599) lower than SA (Table 5). COW PSV decreases by a factor of 0.773 (95% CI [-0.954, −0.613], *p* = 0.00400), and MCA PSV decreases by a factor of 0.955 (95% CI [-1.12, −0.773], *p* = 0.00400) compared to SA (Table 5). COW PDV also decreases by a factor of 0.984 (95% CI [-1.24, −0.721], *p* = 0.00400), and MCA PDV decreases by a factor of 1.01 (95% CI [-1.29, −0.747], *p* = 0.00400) compared to SA (Table 5). Fpr TPVelocity, we were unable to find evidence against the hypothesis that SA = COW = MCA (*p* = 0.861) (COW β = 0.0137, 95% CI [-0.286, 0.313]) (MCA β = 0.0846, 95% CI [-0.224, 0.393]) (Table 5).

For most CSF flow parameters, COW and MCA values for are similar (Table 5). It should be noted that the 95% CI and *p*-value (*p* = 0.0280) for PSV contradict each other, but the latter is more directly related to our hypothesis testing (Table 5). Therefore, we found evidence against the hypothesis that COW = MCA for PSV.

There is considerably greater intra-subject variation than between-subject variation for the LMMs and GLMMs which is unsurprising given the pulsatile nature of CSF flow (Table 5). This result also supports that we found these effects among healthy participants since they are similar to each other.

Sex and age are weak predictors of CSF flow parameters

For all CSF flow parameters, sex, age, and their interaction are on the order of magnitudes less than the intercept (Table 6). Moreover, we did not find evidence against the hypothesis that Sex = 0, Age = 0, and their interaction = 0 (Table 6). While the 95% CIs and *p*-values of SA DFR (*p* = 0.996), COW DFR (*p* = 0.548), MCA DFR (*p* = 0.999), and MCA TPStroVol (*p* = 0.108) contradict each other, we will defer to the latter as explained previously (Table 6). Therefore, we can say with great certainty that the effects of sex, age, and their interaction are weak.

## Discussion

Our results compare favorably with previously reported CSF flow characteristics. We demonstrated pulsatile CSF flow and SA VFR (0.00700 mL/s) in the range of literature values, 0.0049-0.0432 mL/s (Table 2) (Fig. 2) (7, 17–19). However, Florez et al. (16) found SA VFR was 0.0635 mL/s in healthy participants. This difference may be attributed to Florez et al.’s use of background correction and their own semi-automated program for creating ROIs. SA Velocity also fell into the reported range of Stahlberg et al.’s (1989) (20) study: 0-3 cm/s (Table 2). Overall, these results support the use of our semi-automated program.

SA peak velocities were similar and lower compared with the literature. SA PSV (1.45 cm/s) and the magnitude of SA PDV (0.867 cm/s) ranged from 2.0 to 11.5 cm/s in the literature (Table 2) (7, 16, 20, 21). Differences may be attributed to variations in details of the analysis by the use of tolerance in our semi-automated program (Additional file 1) as well as commonly known partial volume effects which cause reduction in maximum flow values, so peak velocities are underestimated.

Generally, differences in CSF flow parameters may also be due to differences in MRI manufacturer, artifacts, resolution levels, and VENC (2).

What is unique to this study though is the examination of CSF flow in the COW and MCA. Since our semi-automated program was validated through the SA results, we can now expand its use to measurements of the COW and MCA. For all of the CSF flow parameters, there is ample evidence that COW and MCA are different than SA (Table 5). Since the magnitude of flow parameters is greater while the magnitude of velocity parameters is lower in the COW and MCA than SA, it is plausible that the size of these conduits is driving the increased flow rates in the COW and MCA compared to SA (Table 5). The area of COW and MCA perivascular space is larger than SA, so this finding is unsurprising. Between the COW and MCA however, there was minimal difference between them for most parameters (Table 5). Since fluid flows through the COW and enters the MCA shortly after, this result was expected.

Besides conduits, demographics may also influence CSF flow dynamics. We used our semi-automated program to establish baseline values of CSF flow parameters for each sex and age group. Moreover, we looked at sex, age, and their interaction as predictors. Across the board though, these effects were negligible. Thus, sex and/or age seems to have minimal influence on CSF flow dynamics. Like our study, Sartoretti et al. (1) used regression models with sex and age as predictors , and they found sex and age could only explain a small part of CSF flow parameters which they quantified to be 6-18%. Furthermore, they found sex and age were not significant predictors for the SA Velocity. Other studies found similar results where sex and age were not significant predictors for the SA VFR, SFR, DFR, Velocity, and Peak Velocity (16, 19, 22, 23).

Some studies, however, have found sex and age dependencies of several CSF flow parameters. Sartoretti et al. (1) found that these predictors were significant for the SA VFR as well as SA peak velocity. The SA VFR and peak velocity increased with age and was higher in males (1). Unal et al. (22) also observed the age dependence of SA peak velocity, but the relationship was inverse. Stoquart-ElSankari found the age dependence for SA VFR, but similarly, the trend was downward. Rohilla et al (8) found weak positive linear correlations with age for the SA SFR and PDV and moderate positive linear correlations for the SA DFR and PSV.This variation in results may be due to the age range of participants and how they were divided into groups. Rohilla et al.’s (8) study had participants from 40 to 78 years of age while our study had participants ranging from 23 to 59 years of age. The elderly group in Stoquart-ElSankari et al.’s (12) study had a mean age of 71 while the young group had a mean age of 27.5. The difference in CSF flow parameters between these groups may be more obvious because of the higher prevalence of chronic conditions among the elderly. Our study, though, only looked at healthy, relatively young participants.

Our results may have also differed because of our limited sample size. However, Sartoretti et al.’s study (1) had 128 healthy participants from 17-88 years of age which found similar results. As both of our studies suggest, other factors may influence CSF flow dynamics to a greater extent, including cardiac pulsations, breathing, anatomy of brain, size of blood vessels, body positioning, exercise, and hydration (1, 7).

Limitations of our study include eddy currents and partial volume effects. Thus, MRI protocols should be optimized, and the effect of different VENCs on CSF flow parameter measurements should be evaluated. Another major limitation is the inaccuracy of the ROI delineation process. Our semi-automated program used for that process could potentially introduce variability in measurements (Additional file 1). Furthermore, it is only capable of capturing one continuous ROI. Thus, if conduits appear in multiple areas of the MRI phase image as in the case of the COW and MCA, measurements are underestimated. To improve the ROI delineation process, our semi-automated program should be formally evaluated, and its inter-observer reliability should be measured (Additional file 1). We believe these efforts would be fruitful as automation has the benefits of increasing accuracy, reproducibility, and the ease of studying large samples (23). Lastly, it is clear there is no consensus on the effects of sex, age, and their interaction, even in literature on the SA. Further studies, then, need to be conducted and particularly focus on comparing the elderly population to younger populations.

In this study, we have established the baseline values for the SA, COW, and MCA parameters as well as highlighted the limited influence of sex and/or age. Future studies can use this research as a starting point to investigate the COW and MCA, thereby increasing the accuracy of parameter measurements. It might also be helpful to look at sex, age, and other factors (e.g., breathing) simultaneously to get a better sense of what drives CSF flow dynamics. Finally, these studies should be repeated in patients with CNS or cerebrovascular system pathologies which could potentially lead to the applications of CSF flow to clinical diagnosis, monitoring, and treatment.

## Notes

### Competing Interest Statement

The authors have declared no competing interest.

### Summary of Updates

Corresponding author updated to reflect co-corresponding authors.

