## Additional file 1 for "Quantitative Evaluation of Normal Cerebrospinal fluid flow in Sylvian aqueduct and perivascular spaces of middle cerebral artery and circle of Willis using 2D phase-contrast MRI imaging"

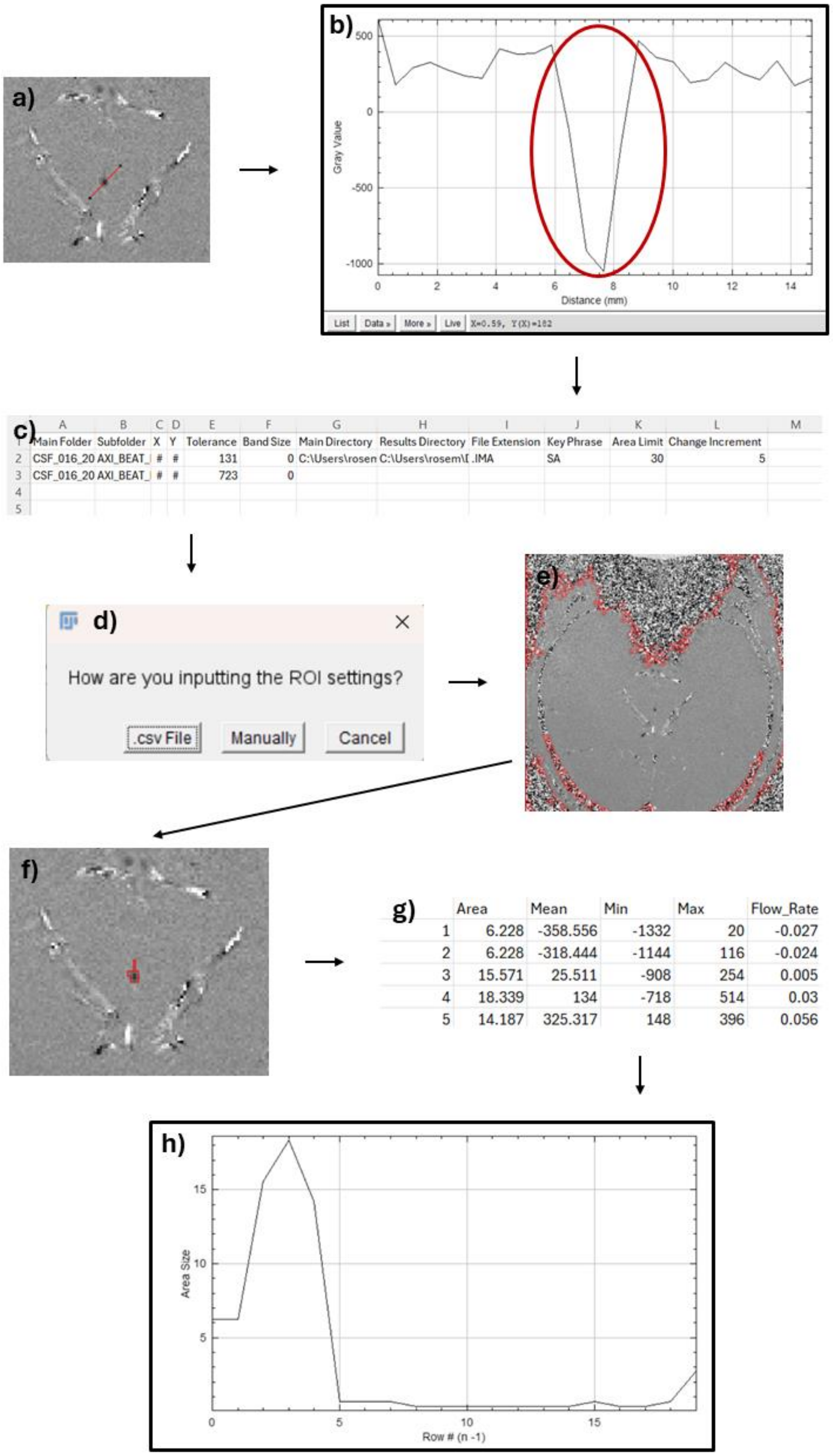

**Process of method with semi-automated program.** a) Identifying the “X-Coordinate” and the “Y-Coordinate.” b) Identifying the peak intensity of the ROI to calculate “Tolerance.” c) Inputting settings for the ROI into a .csv file. d) Methods for users to define settings for ROIs. e) Initial ROI based solely on user-defined settings. f) ROI from “e” after automated program adjustment. g) Results table from program. h) Graph of ROI sizes provided at the end for quality check.

Our semi-automated program was written in the ImageJ/Fiji Macro language. This macro is able to create ROIs from user-defined settings and find flow rate and velocity from those ROIs automatically. There are two ways of defining the settings for ROIs: manually typing them in the program or storing them in a .csv file. The latter lends itself more to bulk analysis, so we used this method for our study.

To figure out the settings for the ROIs, an individual looked at the first phase image of each series. The center of the ROI was found, and a line was marked through it. A plot profile was created by ImageJ/Fiji based on this line. The peak that corresponded to the ROI and its range of intensity values were identified. "Tolerance" was calculated as (the maximum of the peak - the minimum of the peak) / 2. The mode for "Tolerance" was automatically set as "8-connected" in the program. The "X-Coordinate" and "Y-Coordinate" were taken from a point in the ROI that had a similar intensity to the approximate median of the peak's range. "Band Size" was defined as zero for SA and one for COW and MCA. These previously mentioned steps were repeated for each series of images. "Area Limit" was set at 30 for SA and 90 for COW and MCA while "Change Increment" was set at five for all conduits. These values were chosen based on the use of test data with our semi-automated program. All of this information was stored in a .csv file.

The steps of our semi-automated program will now be outlined.

1. Based on the user-defined "Key Phrase," the program parses only through certain folders. For example, if my key phrase was "SA," it would only analyze image series in folders that have "SA" somewhere in its name.
2. The program only opens images that have the user-defined "File-Extension."
3. It finds the settings in the .csv file corresponding to the image series and automatically creates a ROI based on those settings. "X-Coordinate" and "Y-Coordinate" are used to identify the central location of the ROI. "Tolerance" is used to define the range of acceptable intensity values which allows each ROI to be adjusted for a specific image. "Band Size" is used to create a band of the ROI and define the band's size.
4. The program takes initial measurements of the area and mean intensity.
5. If the area exceeds the "Area Limit," the program automatically reduces the "Tolerance" a certain percentage according to the "Change Increment."
6. A new ROI is created based on these new settings.
7. Steps 4-6 are repeated until the area does not exceed the "Area Limit."
8. FR and Velocity are automatically calculated from the area and mean intensity, and all of these values are outputted as a .csv file.
9. A plot of the area vs. each ROI is created for quality check. Possible outliers in area can be identified.

Our semi-automated program also has additional features. Specific ROIs can be checked as needed before running all of the images through the program. Warnings will appear if settings are missing for specific ROIs. ROIs can be given labels.
